# Essential magnetosome proteins MamI and MamL from magnetotactic bacteria interact in mammalian cells

**DOI:** 10.1101/2023.12.19.572379

**Authors:** Qin Sun, Liu Yu, Sarah C. Donnelly, Cecile Fradin, R. Terry Thompson, Frank S. Prato, Donna E. Goldhawk

## Abstract

To detect cellular activities deep within the body using magnetic resonance platforms, magnetosomes are the ideal model of genetically-encoded nanoparticles. These membrane-bound iron biominerals produced by magnetotactic bacteria are highly regulated by approximately 30 genes; however, only a few magnetosome genes are essential and may constitute the root structure upon which biominerals form. To test this, essential magnetosome genes *mamI* and *mamL* were expressed as fluorescent fusion proteins in mammalian cells. Localization and potential protein-protein interaction(s) were investigated using confocal microscopy and fluorescence correlation spectroscopy (FCS). Enhanced green fluorescent protein (EGFP)-MamI and the red fluorescent Tomato-MamL displayed distinct intracellular localization, with net-like and punctate fluorescence, respectively. Remarkably, co-expression revealed co-localization of both fluorescent fusion proteins in the same punctate pattern. An interaction between MamI and MamL was confirmed by co-immunoprecipitation. In addition, changes in EGFP-MamI distribution were accompanied by acquisition of intracellular mobility which all Tomato-MamL structures displayed. Truncation of the MamL C-terminal cationic peptide partially disrupted MamI-MamL colocalization but not mobility. Analysis of extracts from these cells by FCS was consistent with an interaction between fluorescent fusion proteins, including an increase in particle radius. Co-localization and interaction of MamI and MamL demonstrate that these essential magnetosome proteins may have a role in assembly of the magnetosome in any cell type.

## Introduction

As advancements in non-invasive imaging technology continue to shape medical practice (1), there is a pressing need for greater *in vivo* molecular detail, not only to better understand the location and migration of disease processes but also temporal changes in specific gene expression (2). For this type of molecular imaging, endogenous contrast agents offer huge advantages, tracking cellular activities throughout their life cycle and responding to *in vivo* cues. Magnetic resonance imaging (MRI) has the needed soft tissue resolution and depth of penetration in a non-ionizing platform; nevertheless, cellular imaging relies heavily on exogenous contrast agents (3). To expand the capabilities of molecular MRI in mammalian cells, we are using magnetosome genes to program the synthesis of iron particles. Since this form of iron contrast can be genetically regulated, it may be suitable for magnetic resonance (MR) detection of reporter gene expression (4). Magnetotactic bacteria (MTB) synthesize magnetosomes, which consist of an iron biomineral (often magnetite, Fe_3_O_4_) sequestered within a membrane-bound compartment (5, 6). This form of superparamagnetic iron oxide constitutes an ideal MR contrast agent (7, 8) and if mammalian cells could replicate the program of magnetosome synthesis, then MRI at clinical field strength could theoretically detect as few as 3 cells in small animals and approximately 1000 cells in large animals or humans (4).

Magnetosome biosynthesis is under the control of approximately 30 genes, many of which are clustered on a magnetosome genomic island (6, 9, 10). While the proposed functions of these bacterial genes have been divided into vesicle formation and crystal formation (11), most genes are non-essential, as deletion of only a select few genes completely abrogates magnetosome biosynthesis (12). In addition, these essential genes are among those required for vesicle formation in MTB (13), which is the first step in magnetosome formation and ensures a compartment where iron can be safely concentrated for biomineralization, without introducing cytotoxicity. We further propose that essential magnetosome genes constitute the scaffold upon which the fuller magnetosome structure relies (4). This hypothesis implicates select magnetosome proteins in designating where the structure will assemble and therefore anchoring the nanoparticle in a given membrane. If correct, this also suggests that a rudimentary magnetosome-like particle could be genetically programmed in a variety of cell types and for a variety of applications, dependent on the final crystal structure (14).

Several groups have reported MR contrast enhancement using single transgene expression systems like *magA*, encoding a putative iron transporter in species of *Magnetospirillum* (15, 16) or *mms6*, encoding an iron crystallizing protein (17, 18). However, no one iron-handling protein can recreate the full magnetosome structure and most can be individually deleted without compromising the entire structure (12, 19). The number and function of essential magnetosome genes, upon which the entire magnetosome structure relies, is still poorly understood. In general, these genes are widely conserved across species of MTB, are found on the *mamAB* operon, and frequently display preserved gene synteny (6, 19). Moreover, their presence defines the magnetosome as a hierarchical protein structure, containing a common base upon which assembly of the final magnetosome form depends.

Since forming the vesicle that will protect the cell from iron toxicity has been established as a first step in magnetosome synthesis (20), we speculate that less genetic information may be required to initiate magnetosome synthesis in eukaryotes than in prokaryotes, the former of which inherently direct vesicle formation through the Golgi apparatus. In this case, replicating a magnetosome-like particle in eukaryotic cells will only rely on the expression of genes required for designating the compartment, by using specific protein-protein interactions to direct subsequent iron biomineralization. We have tested this hypothesis using *mamI* and *mamL*, two magnetosome genes which are conserved among magnetite-producing MTB (19), are essential to vesicle formation, and when deleted, prevent magnetosome formation (10, 19). We therefore expressed fluorescent fusion proteins of MamI and MamL, alone and together, in a mammalian cell line. Using confocal microscopy and fluorescence correlation spectroscopy (FCS), we then demonstrated the nature of their intracellular membrane localization in a foreign cell environment and provided evidence of their co-localization and potential for interaction. These results clarify our understanding of the foundation for magnetosome assembly and highlight previously unrecognized activities deemed essential to support iron nanoparticle biosynthesis.

## Results

### Magnetosome gene expression in a mammalian system

Like other MTB (21–23) and magnetosome (18) proteins expressed in mammalian cell lines, MamI and/or MamL were stable in long-term cell culture, with little or no effect on cell viability. Cells were transfected with pEGFP/*mamI* and ptdTomato/*mamL* constructs, either alone or in combination, to obtain stable expression of N-terminal fluorescent fusion proteins. Compared to cells expressing EGFP alone, western blots revealed an increase in the size of α-EGFP immuno-stained bands in samples from EGFP-MamI-expressing cells (Fig. 1, A and C). The size shift was consistent with the expected molecular weight (MW) of MamI (approximately 8 KDa). Similarly, compared to cells expressing Tomato from the empty vector, western blots revealed an increase in the size of α-tdTomato immuno-stained bands in samples from Tomato-MamL-expressing cells (Fig. 1, B and C). This size shift was likewise commensurate with the reported MW of MamL (approximately 13 KDa).

**Figure 1.**
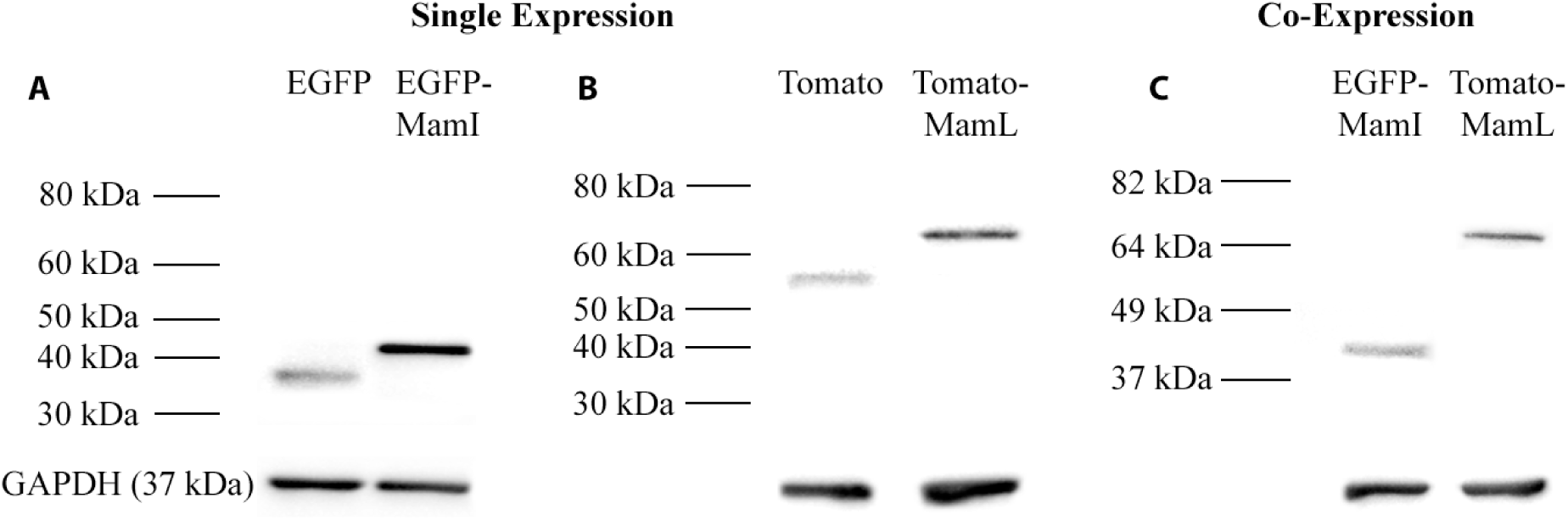
Immunoblots of mammalian cells expressing fluorescent magnetosome fusion proteins. Total cellular protein from MDA-MB-435 cells stably expressing either EGFP or EGFP-MamI (*A*), Tomato or Tomato-MamL (*B*), or both magnetosome fusion proteins (*C*) was examined by western blot, using mouse α-EGFP (*A*, *C*) and/or rabbit α-Tomato (*B*, *C*) as the primary antibodies. Type of fluorescent protein expressed by the cells is indicated above each lane. In panel C, the same cell sample was probed for each magnetosome fusion protein. Approximate MW is shown in the left margin. The loading control was GAPDH (bottom panels). Full-length blots are presented in Fig. S1.

The pattern of intracellular fluorescence was examined in living cells using confocal fluorescence microscopy. Cells expressing EGFP-MamI produced a net-like pattern of green fluorescence (Fig. 2), distinct from the diffuse fluorescence of EGFP alone (Fig. 2, A and B). The atypical pattern of EGFP-MamI fluorescence neither circumscribes the plasma membrane nor outlines the standard shape of intracellular vesicle (Fig. 2, C and D). In contrast, cells expressing Tomato-MamL displayed punctate red fluorescence throughout the cell (Fig. 3). Unlike the uniformly diffuse fluorescence of Tomato alone (Fig. 3, A and B), Tomato-MamL fusion protein appeared in discrete intracellular points, with little or no labelling of the plasma membrane (Fig. 3, C and D).

**Figure 2.**
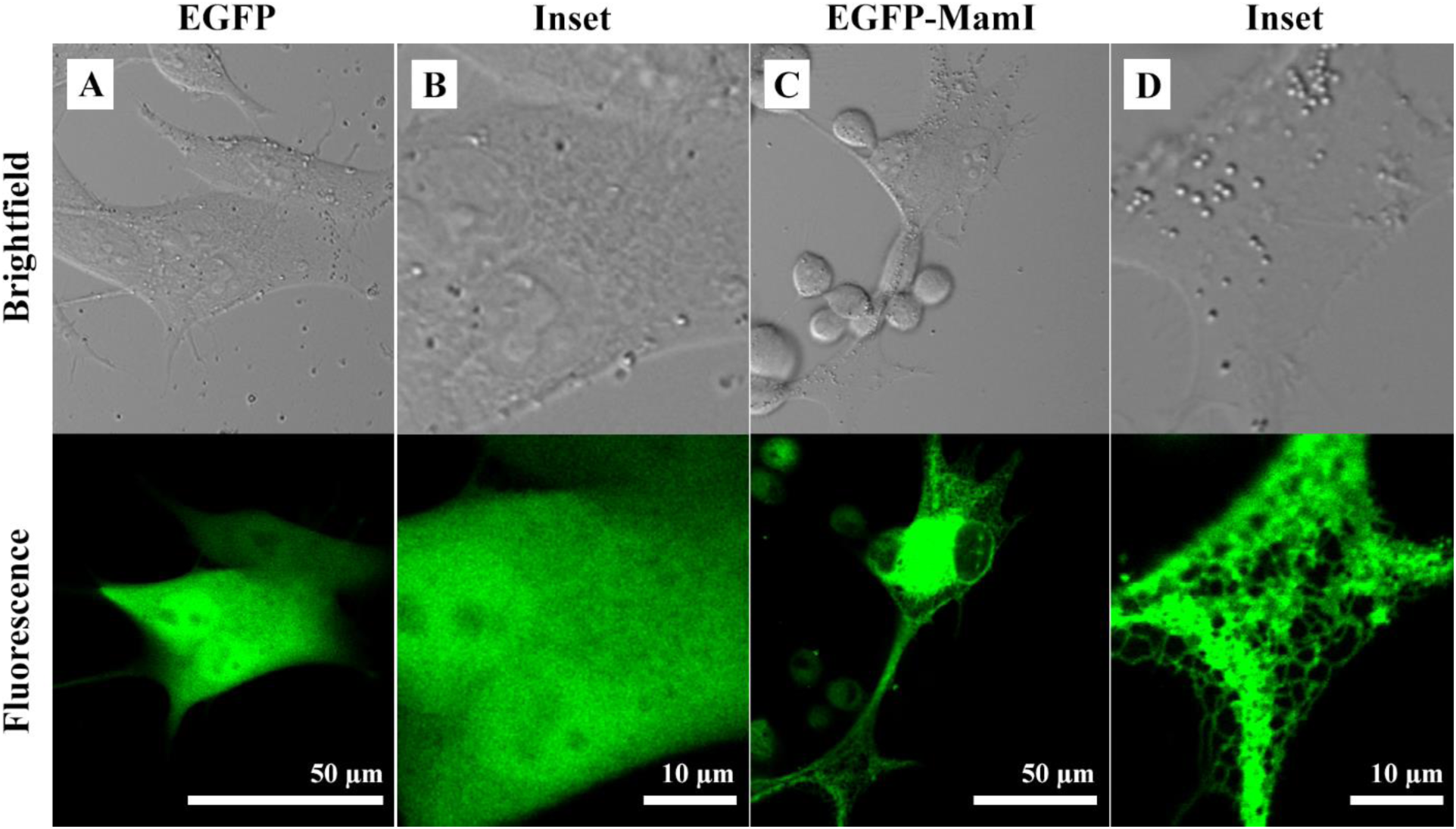
Confocal fluorescence microscopy of mammalian cells stably expressing EGFP-MamI fusion protein. Transfected cells were placed under selection and enriched using FACS to obtain populations expressing EGFP alone (*A* and *B*) or fused to MamI (*C* and *D*). Compared to the uniform fluorescence pattern of EGFP in the cytosol, EGFP-MamI fusion protein displays a net-like pattern of intracellular fluorescence.

**Figure 3.**
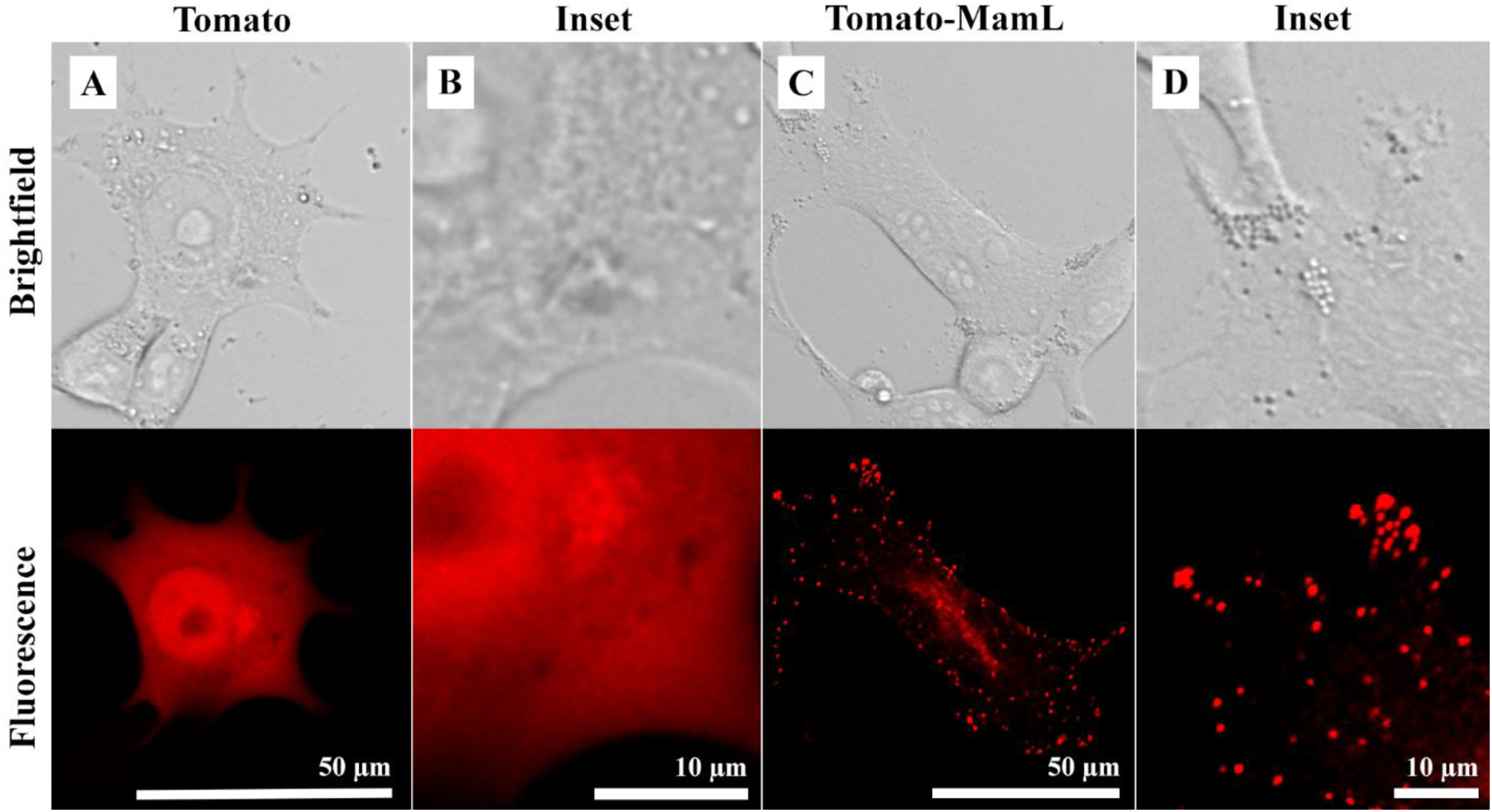
Confocal fluorescence microscopy of mammalian cells stably expressing Tomato-MamL fusion protein. Transfected cells were placed under selection and enriched using FACS to obtain populations expressing Tomato alone (*A* and *B*) or fused to MamL (*C* and *D*). Compared to the uniform fluorescence pattern of Tomato in the cytosol, Tomato-MamL fusion protein displays a punctate intracellular fluorescence pattern. These punctate structures are dispersed throughout the cell and mobile (Movie 1B).

### Mobility of MamL in mammalian cells

Interestingly, Tomato-MamL expression in mammalian cells produced mobile structures (Movie 1, GIF attachment). Compared to the green fluorescent structures of EGFP-MamI, whose location remained relatively stable (Movie 1, A arrow), some of the Tomato-MamL red fluorescent particles exhibited considerable displacement (Movie 1, B arrow). In addition to movement within the x-y plane of focus, the pattern of red fluorescence appeared to dip in and out of the focal plane, indicating potential movement of Tomato-MamL along the z-axis.

### Mobility of truncated MamL

Structural modeling of MamL predicts that a cationic C-terminal peptide lies outside the transmembrane domain (12). To assess the influence of this putative extramembrane domain on Tomato-MamL localization and mobility, the C-terminal 15 amino acids were removed, permitting expression of a truncated Tomato-MamL fusion protein. Western blotting confirmed the size difference between full-length (67 kDa) and truncated (65 kDa) MamL (MamL_trunc_, Fig. 4). Confocal fluorescence microscopy confirmed the presence of mobile, punctate Tomato-MamL_trunc_ particles in mammalian cells (Fig. 5A and Movie 2, GIF attachment). However, over half of the cell population also displayed a diffuse pattern of red fluorescence that was stationary (Fig. 5B).

**Figure 4.**
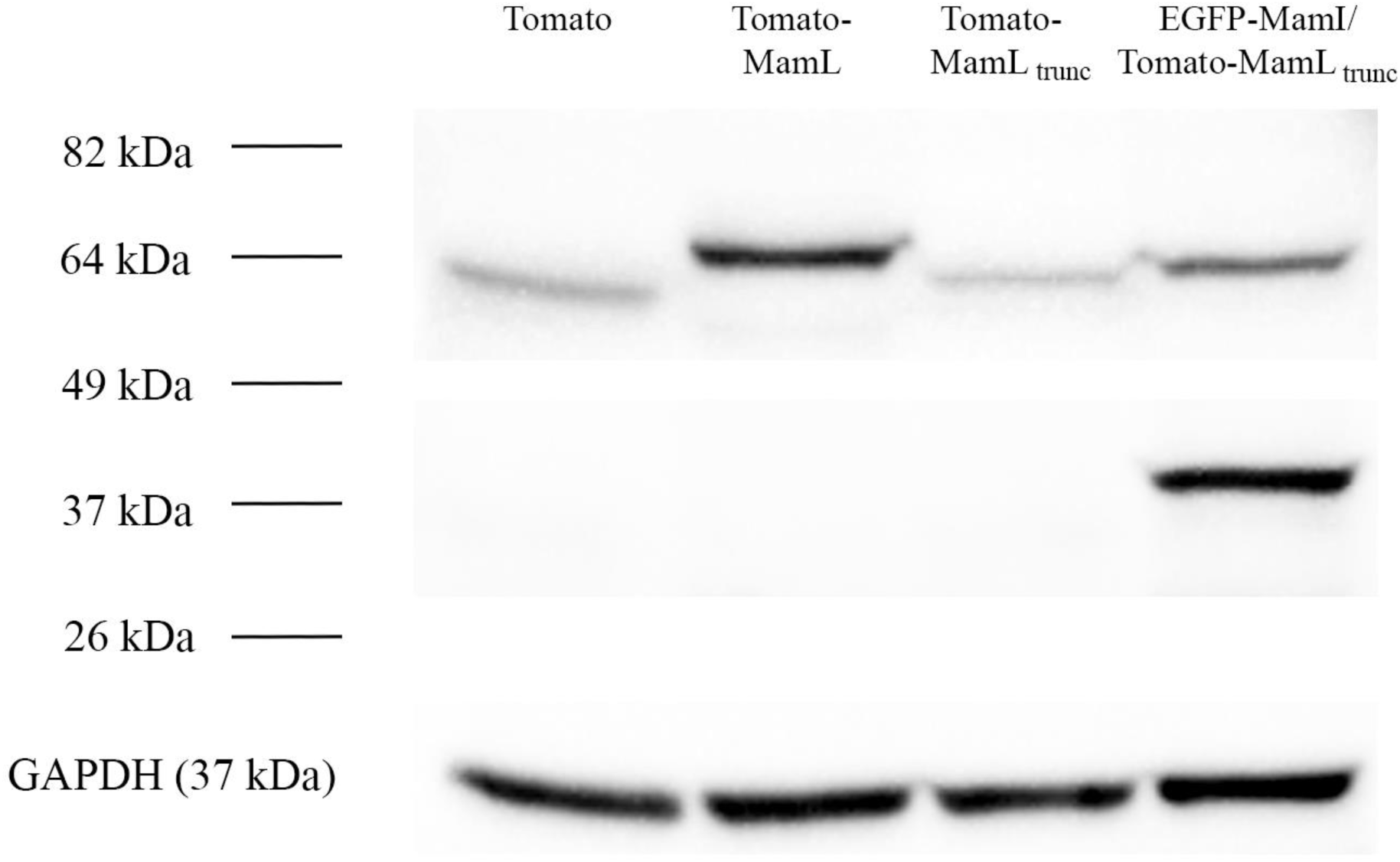
Expression of fluorescent magnetosome fusion proteins in mammalian cells. Total cellular protein from MDA-MB-435 cells stably expressing either Tomato, Tomato-MamL, Tomato-MamL_trunc_, or both EGFP-MamI and Tomato-MamL_trunc_ was examined by western blot. Type of fluorescent protein expressed by the cells is indicated above each lane. For EGFP-MamI/Tomato-MamL_trunc_ extracts, the same cell sample was probed for each magnetosome fusion protein. Approximate MW is shown in the left margin. The loading control was GAPDH (bottom panel). Full-length blots are presented in Fig. S2.

**Figure 5.**
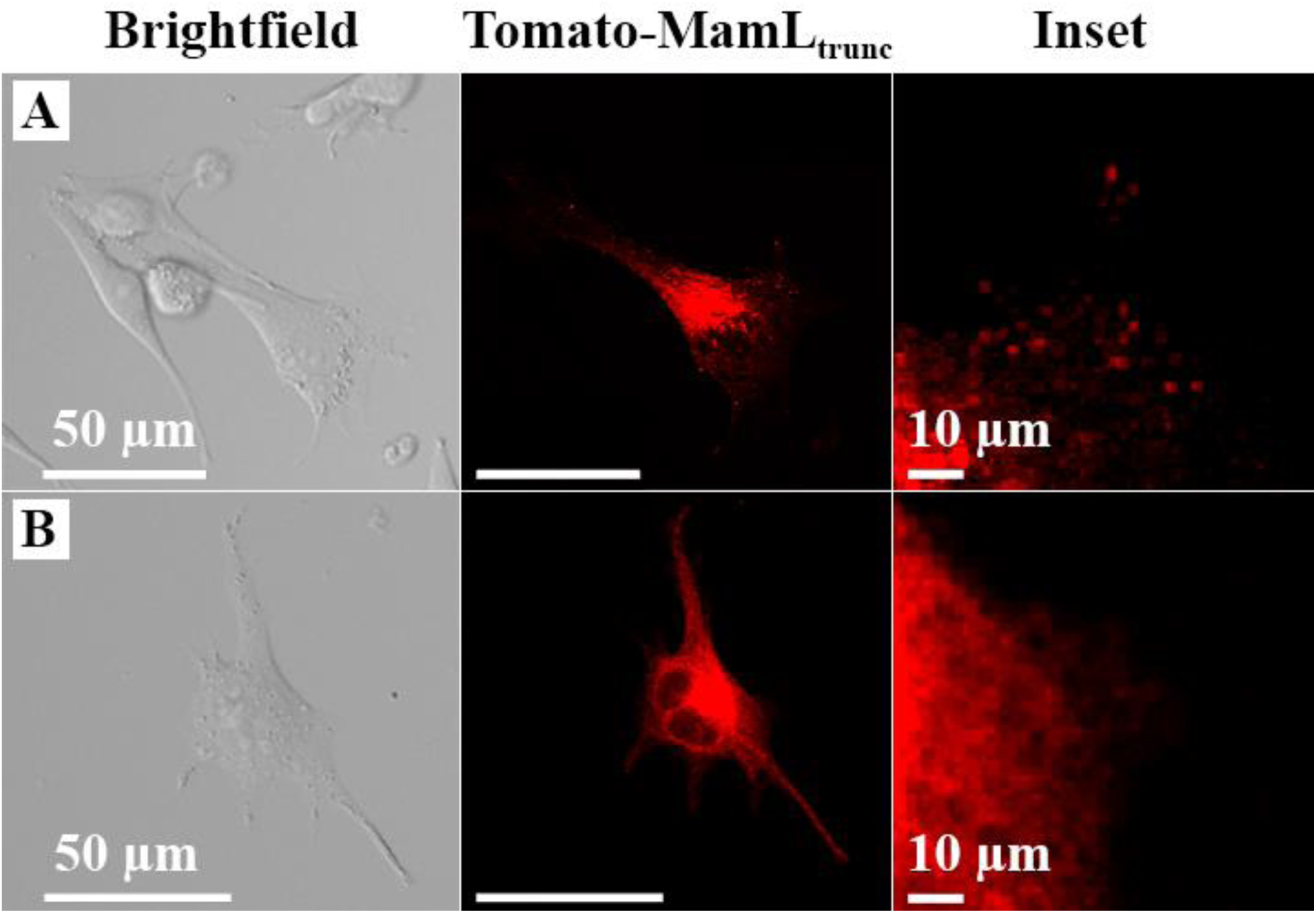
Confocal fluorescence microscopy of mammalian cells stably expressing Tomato-MamL_trunc_ fusion protein. The population of cells expressing Tomato-MamL_trunc_ displayed a punctate pattern (A) in approximately 40% of cells and a diffuse pattern (B) in the remaining 60% of cells. Punctate structures were dispersed throughout the cell and mobile, similar to full-length Tomato-MamL (Movie 2).

### Co-localization of MamI and MamL

Stable co-expression of EGFP-MamI and full-length Tomato-MamL yielded a population of cells that was enriched for both red and green fluorescence by FACS (Fig. 6). When co-expressed with Tomato-MamL, the pattern of EGFP-MamI fluorescence was no longer net-like. Instead, green fluorescence was now punctate (Fig. 6, B), similar to the pattern of Tomato-MamL (Fig. 6, C). Moreover, when both red and green channels were superimposed, yellow punctate fluorescence was observed throughout the cells (Fig. 6, D), indicating co-localization of EGFP-MamI and Tomato-MamL.

**Figure 6.**
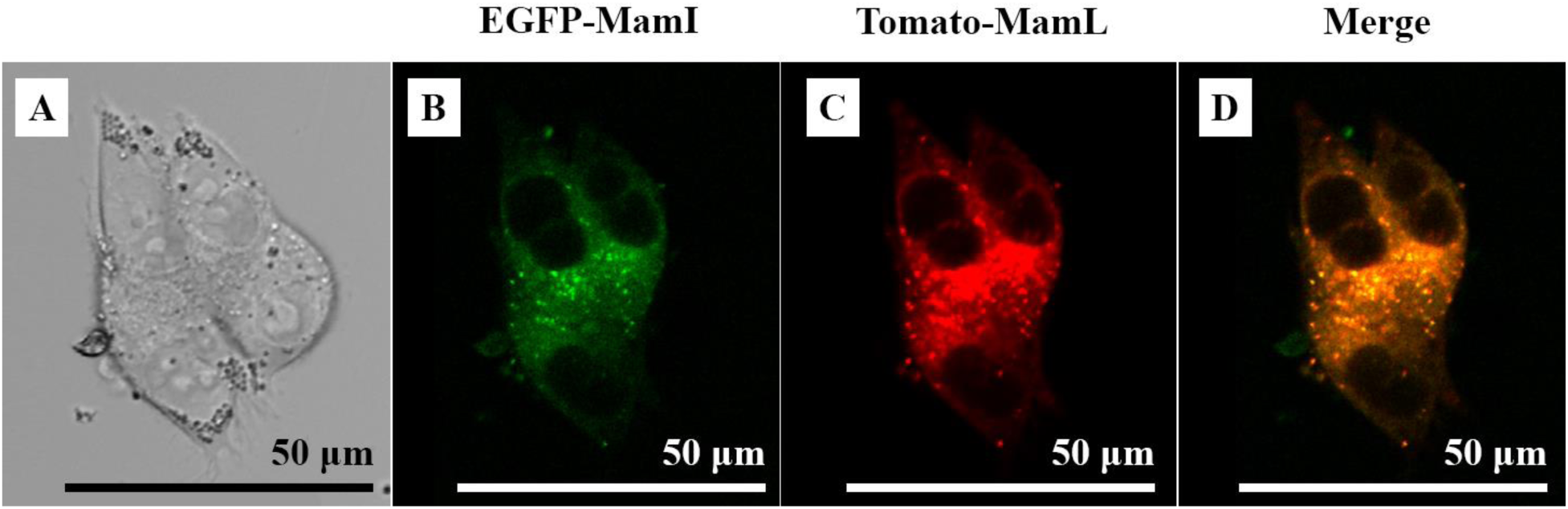
Confocal fluorescence microscopy of mammalian cells stably co-expressing Tomato-MamL and EGFP-MamI fusion proteins. Transfected cells were placed under selection and enriched using FACS to obtain a population co-expressing Tomato-MamL and EGFP-MamI. Dividing cells shown in brightfield (*A*) are displayed under green (*B*) and red (*C*) fluorescence, highlighting the expression of EGFP-MamI and Tomato-MamL, respectively. Merging green and red fluorescence (*D*) indicates colocalization of fusion proteins in a yellow, punctate intracellular fluorescent pattern. These punctate structures are dispersed throughout the cell and mobile (Movie 3).

Remarkably, the same pattern of mobility was displayed by both EGFP-MamI and Tomato-MamL when co-expressed in the same cells. This is illustrated in three separate cines obtained from either the green fluorescent channel, the red fluorescent channel, or both channels simultaneously (Movie 3, GIF attachment). In each channel, arrows point to punctate structures as they move in and out of the plane of focus and laterally within the focal plane. Whether tracking particles containing EGFP-MamI (Movie 3, A), Tomato-MamL (Movie 3, B) or both fusion proteins (as identified by a yellow color in Movie 3, C), any single group of cells displayed the same punctate pattern of fluorescence and movement in the x-y and z directions.

### Effect of MamL truncation on co-localization with MamI

Compared to full-length MamL, cells expressing Tomato-MamL_trunc_ displayed more diffuse (∼60%) than punctate (∼ 40%) red fluorescence (Fig. 5). When MamL_trunc_ is co-expressed with EGFP-MamI, three different patterns of fluorescence are observed: diffuse green and red (∼45%, Fig. 7A), diffuse green and punctate red (∼45%, Fig. 7B), and punctate green and red (10%, Fig. 7C). Removal of the MamL C-terminal peptide partially disrupted co-localization and interaction of the two proteins. When those cells co-expressing MamI and MamL_trunc_ display both green and red punctate fluorescence (Fig. 7C), the co-localization and mobility of these structures is similar to that observed with full-length fusion proteins (Movie 4, GIF attachment). That so few cells display evidence of punctate mobile yellow fluorescence implicates the C-terminal MamL peptide in mediating an efficient interaction with MamI. This is further supported by the presence of many cells with punctate mobile red fluorescence (from Tomato-MamL_trunc_; Movie 5, GIF attachment) in the presence of diffuse green fluorescence (from EGFP-MamI, Fig. 7B).

**Figure 7.**
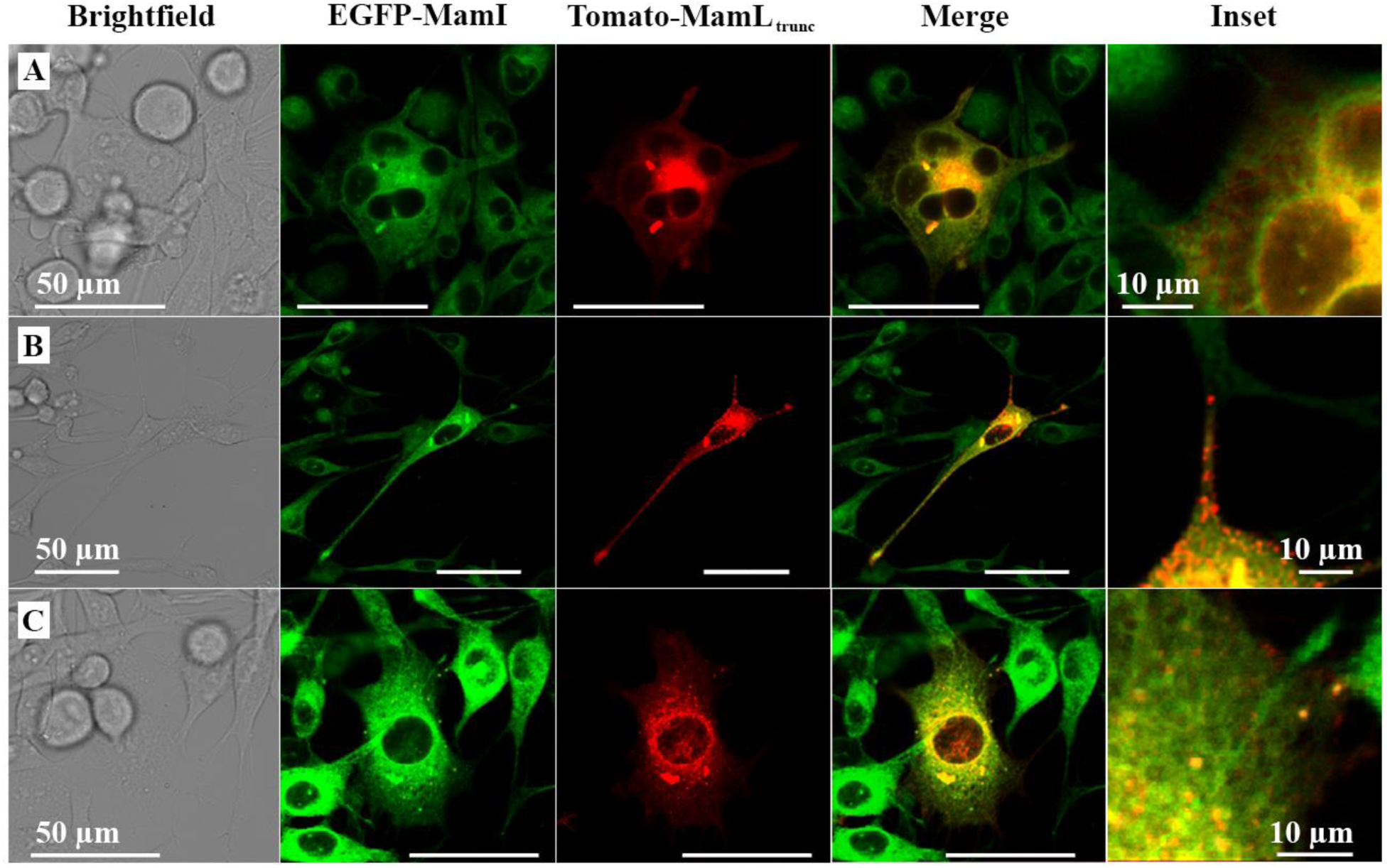
Confocal fluorescence microscopy of mammalian cells co-expressing Tomato-MamL_trunc_ and EGFP-MamI fusion proteins. Cells shown in brightfield are displayed under green (EGFP-MamI) and red (Tomato-MamL_trunc_) fluorescence. Merging green and red fluorescence (Merge) indicates colocalization of fusion proteins in yellow. The inset reveals 3 types of fluorescent pattern in cells co-expressing EGFP-MamI/Tomato-MamL_trunc_: diffuse green and red fluorescence (A), diffuse green and punctate red fluorescence (B), and punctate green and red fluorescence (C). All punctate structures are dispersed throughout the cell and mobile (Movies 4 and 5). Scale bars indicate 50 µm unless otherwise noted.

### Co-localization of FLAG-MamL and EGFP-MamI

To refine the co-expression of MamI and MamL; examine the influence of a bulky fluorescent tag; and facilitate downstream fluorescence analyses of additional magnetosome proteins, we expressed FLAG-MamL in mammalian cells. When co-expressed with EGFP-MamI, the net-like green fluorescent pattern (displayed by EGFP-MamI alone) changes to a mobile, green fluorescent, punctate pattern (Fig. 8, Movie 6, GIF attachment), consistent with an interaction between these magnetosome proteins and little or no interference from the fluorescent moiety. Western blots (Fig. 9) confirm the presence of both EGFP-MamI and FLAG-MamL.

**Figure 8.**
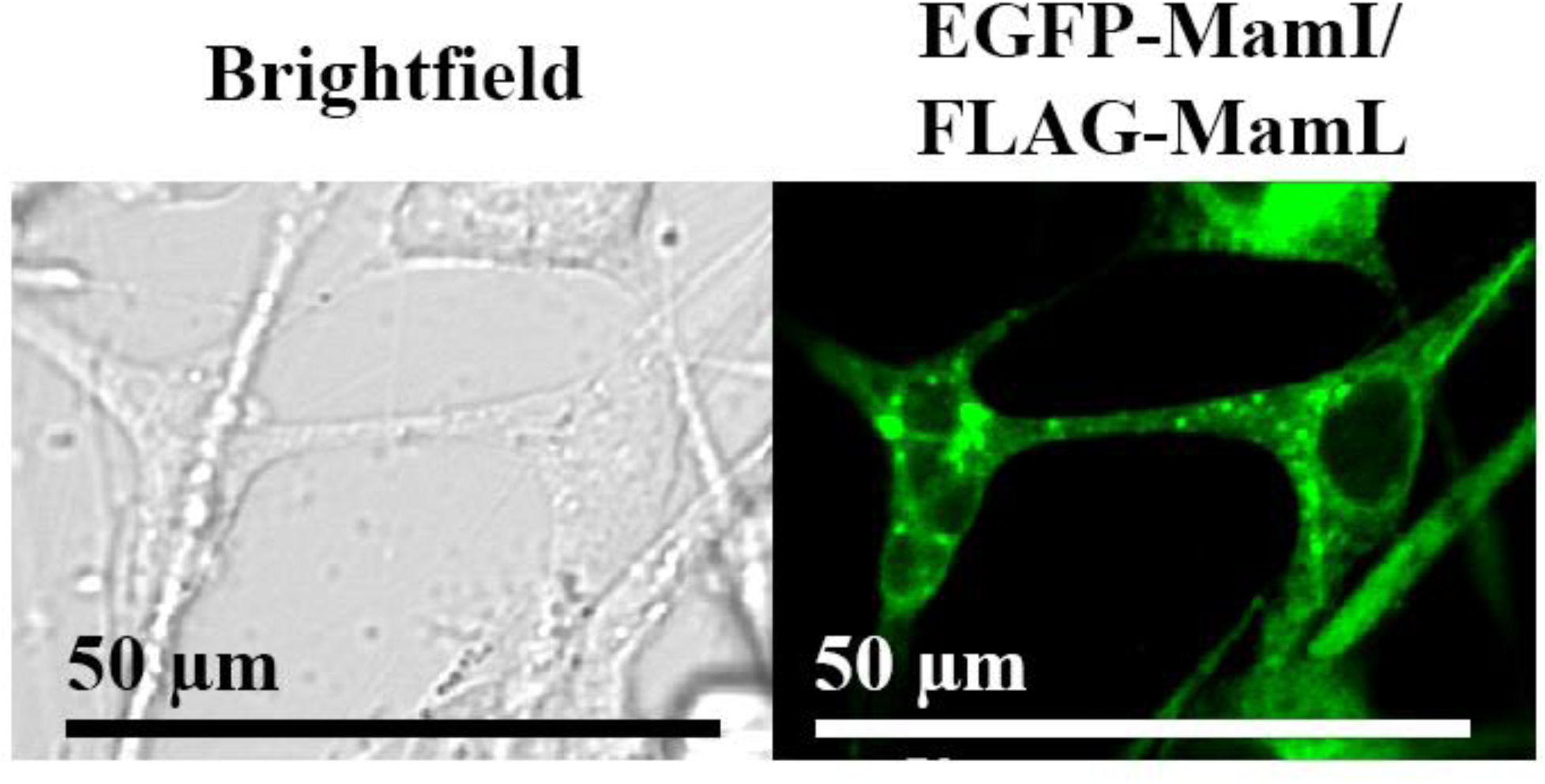
Confocal fluorescence microscopy of mammalian cells stably co-expressing FLAG-MamL and EGFP-MamI. Micrographs display a population of MDA-MB-435 cells co-expressing FLAG-tagged MamL and EGFP-MamI fusion protein. The punctate intracellular fluorescence revealed by EGFP-MamI is similar to that obtained with two fluorescent fusion proteins (Tomato-MamL and EGFP-MamI, Fig. 6). Mobile punctate structures are dispersed throughout the cell (Movie 6).

**Figure 9.**
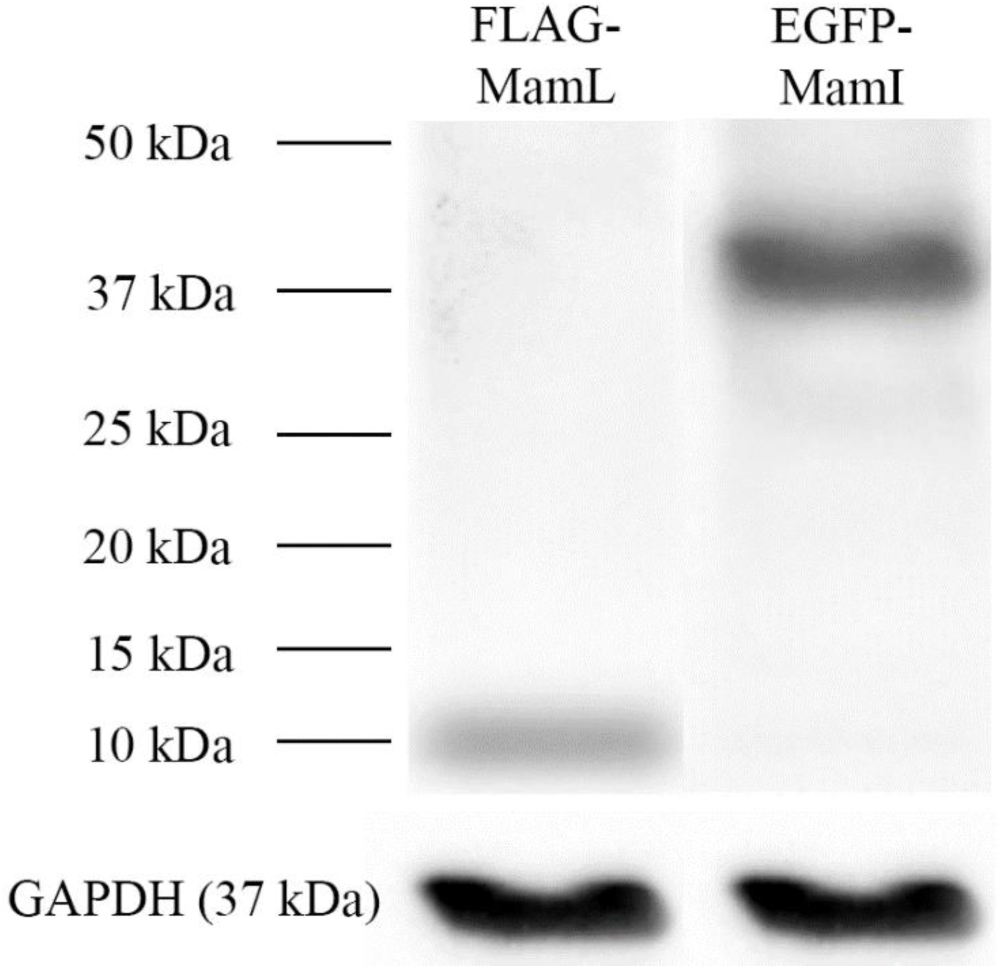
Immunoblots of mammalian cells co-expressing magnetosome proteins. Total cellular protein from MDA-MB-435 cells stably expressing both FLAG-MamL and EGFP-MamI was examined by western blot using primary α-FLAG and α-EGFP antibodies. The same sample was probed in both lanes. Approximate size of FLAG-MamL is 11 kDa. Approximate MW is shown in the left margin. The loading control was GAPDH (bottom panels). Full-length blots are presented in Fig. S4.

### Interactions between MamI and MamL

Apparent MamI-MamL interactions were further examined by FCS in lysed cell samples containing total cellular protein (Fig. S3). Using this technique, the diffusion coefficient of fluorescent species present in the sample was measured and their apparent radius (*R*) was calculated. The diffusion of EGFP-MamI (in the absence of Tomato-MamL) was not significantly different from that of EGFP alone (*D =* 90 μm^2^/s ± 3% vs 108 μm^2^/s ± 4%), as might be expected if EGFP-MamI exists as a monomer or dimer in the cell extract (Table 1). The specific brightness of the EGFP-MamI species detected (10.5 x 10^3^ photon/s) was approximately double that of EGFP (4.9 x 10^3^ photon/s), further suggesting that the fusion protein might be a dimer. On the other hand, in extracts from cells co-expressing both proteins, EGFP-MamI diffused significantly slower than when this fusion protein was expressed alone (*D =* 43.2 μm^2^/s ± 50% vs 90 μm^2^/s ± 3%, p < 0.01), as expected of a larger particle (Table 1). The intrinsic brightness of the fusion protein in the context of co-expression could not be determined in this case, because of heterogeneity in the population of detected particles. The apparent radius of EGFP-MamI when co-expressed with Tomato-MamL (4.4 nm, calculated from the value of the measured diffusion coefficient) was significantly larger (*p* < 0.05) than when EGFP-MamI was expressed alone (2.1 nm).

**Table 1.** FCS parameters in mammalian cell extracts of EGFP, EGFP-MamI or EGFP-MamI/Tomato-MamL.

|  | **EGFP** | **EGFP-MamI** | **EGFP-MamI /**<br>**Tomato-MamL** |
| --- | --- | --- | --- |
| Diffusion coefficient <sup>*</sup><br>($\mu\text{m}^2/\text{s}$) | $108 \pm 4\%$ <sup>†</sup> | $90 \pm 3\%$ <sup>‡</sup> | $43.2 \pm 50\%$ <sup>†,‡</sup> |
| Apparent radius <sup>*</sup><br>(nm) | $1.7 \pm 4\%$ <sup>§</sup> | $2.1 \pm 3\%$ <sup>¶</sup> | $4.4 \pm 50\%$ <sup>§,¶</sup> |
| Intrinsic brightness<br>($10^3$ photon/s) | $4.9 \pm 1\%$ <sup>#</sup> | $10.5 \pm 1\%$ <sup>#</sup> | ND |

<sup>\*</sup> Data are the median +/- standard deviation expressed as a percentage of the median (n = 5).
<sup>†,‡,#</sup> $p < 0.001$
<sup>§,¶</sup> $p < 0.05$
ND, not detectable

In the red channel, a similar pattern was observed for the diffusion of Tomato-MamL (Table 2). Its diffusion coefficient (49 μm^2^/s ± 3%) is very close to that of Tomato (51 μm^2^/s ± 56%), as expected if Tomato-MamL in the cellular extract exists as a monomer. This is confirmed by the very similar intrinsic brightness of Tomato-MamL and Tomato (2.5 x 10^3^ vs. 2.6 x 10^3^ photons/s, respectively). However, in extracts of cells co-expressing Tomato-MamL and EGFP-MamI, the particle is diffusing more slowly (D = 29 μm^2^/s ± 38%) than either Tomato or Tomato-MamL alone, indicating that a larger structure is formed upon co-expression of these fluorescent magnetosome fusion proteins (again, with population heterogeneity precluding the measurement of intrinsic brightness). Based on the value of its diffusion coefficient, the average apparent radius of the Tomato-MamL/EGFP-MamI particle is 6.6 nm (Table 2), comparable to the size measured in the green fluorescence channel (4.4. nm, Table 1).

**Table 2.**
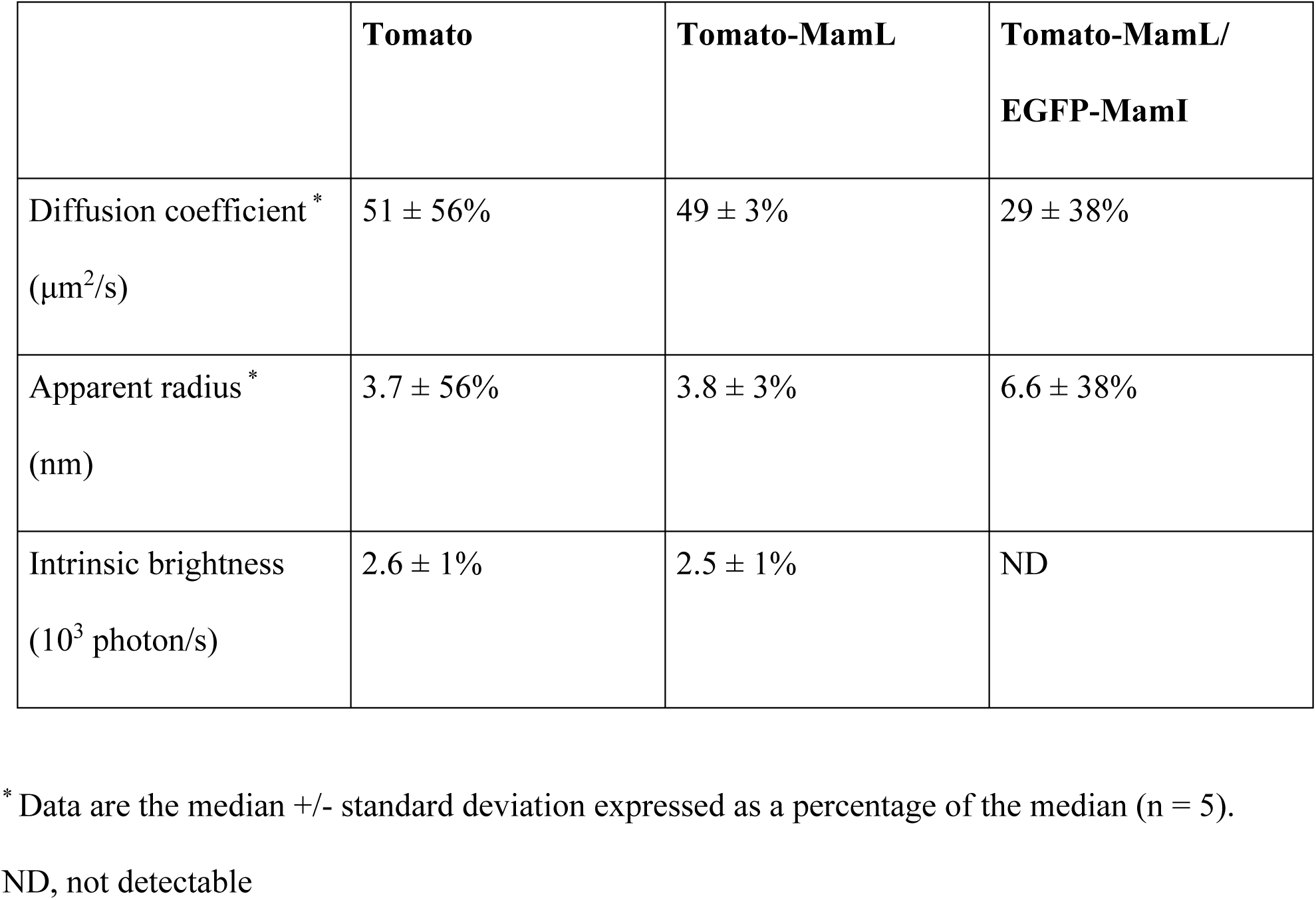
FCS parameters of mammalian cell extracts expressing Tomato, Tomato-MamL, or Tomato-MamL/EGFP-MamI.

|  | **Tomato** | **Tomato-MamL** | **Tomato-MamL/**<br>**EGFP-MamI** |
| --- | --- | --- | --- |
| Diffusion coefficient *<br>($\mu\text{m}^2/\text{s}$) | $51 \pm 56\%$ | $49 \pm 3\%$ | $29 \pm 38\%$ |
| Apparent radius *<br>(nm) | $3.7 \pm 56\%$ | $3.8 \pm 3\%$ | $6.6 \pm 38\%$ |
| Intrinsic brightness<br>($10^3$ photon/s) | $2.6 \pm 1\%$ | $2.5 \pm 1\%$ | ND |

\* Data are the median +/- standard deviation expressed as a percentage of the median (n = 5).
ND, not detectable

Protein-protein interactions between MamI and MamL were further examined by co-immunoprecipitation (Fig. 10). In samples co-expressing EGFP-MamI and FLAG-MamL, both species are detected in the protein pellet after immunoprecipitation with a GFP antibody. The co-immunoprecipitation result is thus consistent with fluorescence microscopy and correlation spectroscopy, and provides further support for the interaction between EGFP-MamI and FLAG-MamL.

**Figure 10.**
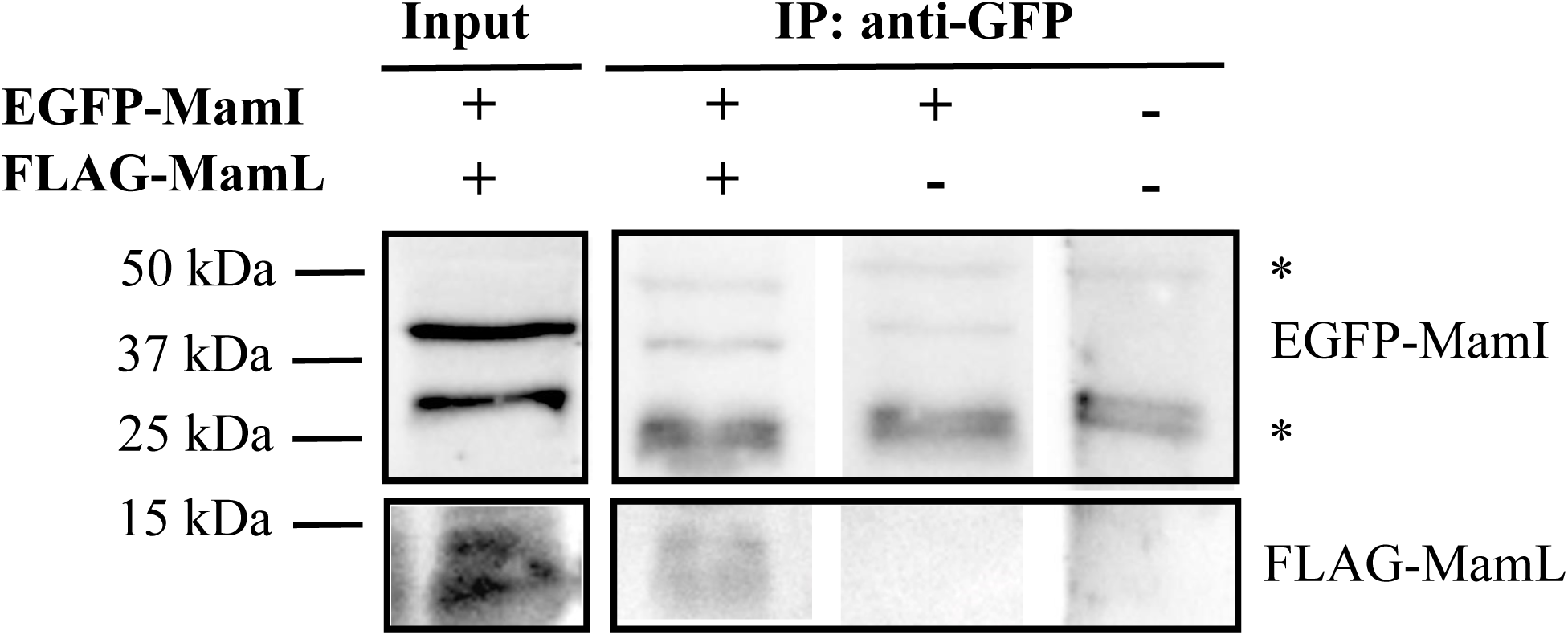
FLAG-MamL co-immunoprecipitates with EGFP-MamI. A western blot of total cellular protein (Input) confirms that cells expressing both EGFP-MamI and FLAG-MamL were used for co-immunoprecipitation (IP) with mouse anti-GFP. Approximate MW is shown in the left margin. Upper panels were probed for EGFP-MamI (∼41 kDa) and lower panels were probed for FLAG-MamL (∼11 kDa). Asterisks indicate immunoglobulin heavy and light chains present in all IP samples. FLAG tagged protein was only detected in samples co-expressing both magnetosome proteins. No FLAG-MamL was detected in IP samples containing total cellular protein from the parental control or those expressing EGFP-MamI alone.

## Discussion

This report describes the expression of fluorescently tagged, magnetosome-associated membrane proteins MamI and MamL in a mammalian cell system and demonstrates their ability to co-localize and interact in a foreign environment. The specificity of this magnetosome protein interaction was characterized by fluorescence confocal microscopy and correlation spectroscopy, showing co-localization of fluorescent fusion proteins, specific recruitment of MamI to MamL particles, as well as interactions between MamL and mobile element(s) within the mammalian intracellular compartment. The bifunctional nature of MamL in intact cells, as revealed by the recruitment of MamI to mobile MamL structures, was verified in cell extracts of co-expressed protein. A decrease in molecular diffusion coefficients was consistent with the formation of species of increasing macromolecular radius, as expected from the interaction of MamI and MamL and confirmed by co-immunoprecipitation.

### Combined magnetosome gene expression in mammalian cells

When co-expressed, the proximity of green and red fluorescence from MamI and MamL fusion proteins, respectively, resulted in yellow particles. This evidence of co-localization is strengthened by changes in a) subcellular localization of MamI, b) diffusion coefficient and apparent radius of co-expressed fusion proteins, and c) co-immunoprecipitation of magnetosome fusion proteins.

Confocal microscopy of live cells expressing EGFP-MamI alone reveals a net-like pattern of green fluorescence, while Tomato-MamL exhibits punctate, red fluorescent structures in the presence and absence of EGFP-MamI. The net-like structure of MamI alone may be related to dimerization of the EGFP fusion protein, whose intrinsic brightness in cellular extracts was twice that of EGFP (10,500 vs 4,500 photons/s). In contrast, Tomato-MamL appears to exist as a monomer in cellular extracts, with intrinsic brightness that is only modestly quenched compared to Tomato (2,500 vs 2,600 photons/s). Although intrinsic brightness values were obtained from FCS measurements in cell extracts (*i.e.* actual cellular structures not preserved), these measurements still give information on the elementary units of protein structures.

The characteristics of singly expressed MamI may be related to the endoplasmic reticulum (ER) which displays a similar net-like pattern in peripheral ER tubules (24). Although biological relevance of this structure within the context of magnetosome assembly is unclear, potentially the interaction between MamI and MamL has features shared with ER-shaping proteins. Super resolution microscopy techniques may shed light on this and the potential role of MamI and MamL in membrane invagination, implicated in magnetosome vesicle formation in MTB (20) and caveolae formation in mammalian cells. Regardless, the net-like pattern of MamI was replaced by punctate green fluorescence in the presence of full-length MamL, whether or not fused to Tomato. Moreover, there was no residual net-like pattern, suggesting that the association of MamI and MamL is the dominant interaction, producing an intracellular structure as expected during initiation of a magnetosome-like particle in a mammalian cell system.

The apparent diffusion coefficient observed for fluorescent MamI and MamL fusion protein structures in mammalian cells, D = 2.2 µm^2^/s, corresponds to that of a particle with a hydrodynamic radius of about 20 nm (assuming a cellular viscosity 5 times that of water). This size is comparable to that of transport vesicles in eukaryotic cells (radius 15 to 50 nm) and to that of magnetosome crystals (average radius 21 nm in *M. magneticum* AMB-1 (25) and between 17 to 60 nm (26) for other MTB species). While there is no expectation of biomineral formation in the present work, in MTB where the full complement of magnetosome genes is expressed, divergent roles for MamI have been reported in different species of *Magnetospirillum*, suggesting that cellular milieu may influence function. While MamI is essential to vesicle formation in *M. magneticum*, the same role has not been assigned in *M. gryphiswaldense* (27). In the latter MTB, a role for MamI in magnetite nucleation has been reported (28). For future applications in molecular MRI, both roles of MamI will be important and may yet be validated as mammalian expression systems are developed to incorporate additional, essential magnetosome genes that optimize the rudimentary magnetosome-like nanoparticle (29).

MamI and MamL are small integral membrane proteins (77 and 123 amino acids, respectively) with predicted alpha helices that span the (magnetosome) membrane twice (12). In extracts where cellular structure is removed, correlation spectroscopy indicated that particles expressing both green and red fluorescence (co-expression of MamI and MamL) diffused more slowly than either fluorescent protein (EGFP or Tomato) or fluorescent fusion protein (EGFP-MamI or Tomato-MamL) alone. As expected, if these two fusion proteins directly interact or localize to the same structure, regardless of which fluorescent channel is used to generate autocorrelation curves (Fig. S3), the diffusion coefficients of co-expressed fusion proteins are similar (approximately 30 to 40 µm^2^/s for red and green fluorescence, respectively). From these measurements, the radius of the structure to which MamI and MamL belong in the extract was estimated to be on the order of 4 to 7 nm. Future work using fluorescence cross correlation spectroscopy will confirm whether or not these fusion proteins are indeed moving together, as strongly suggested by co-immunoprecipitation and changes in their subcellular localization and mobility (discussed below).

### MamL mobility and association with MamI

The mobility of MamL particles in mammalian cells is an unexpected finding, offering new insights into magnetosome substructure and the potential roles of individual magnetosome genes. The presence of yellow fluorescent particles not only confirms the association between EGFP-MamI and Tomato-MamL, but also distinguishes this interaction from that occurring between MamL and cellular mobile elements. If MamI binds directly to MamL, then the interaction between these magnetosome proteins may occur primarily in the membrane compartment, leaving the ionic C-terminal residues of MamL available for alternative functions. However, as shown in this study, the C-terminal peptide of MamL contributes to the co-localization and interaction of MamI and MamL.

### Truncation of the MamL C-terminal peptide

Whether expressed alone or in conjunction with EGFP-MamI, Tomato-MamL structures were mobile. Analysis of the amino acid sequence of MamL reveals a C-terminal peptide of 15 amino acids that is rich in positively-charged amino acids and bears sequence homology to cell-penetrating peptides (CPP) (11). This class of cationic peptides can traverse the plasma membrane by interacting with negatively charged phospholipids (30). Although any such role for MamL awaits further characterization, here we describe the mobility of Tomato-MamL structures in terms of plausible interactions between MamL C-terminal residues and anionic components of the cytoskeleton. This is consistent with structure predictions of MamL, suggesting that its C-terminal cationic residues lie outside the membrane, theoretically exposed to the cytosolic face of an intracellular vesicle (12). CPP have also been reported to interact with cytoskeletal components; for example, the synthetic arginine-rich peptide R_8_W strongly interacts with the anionic proteins actin and tubulin (30, 31). Our analysis of confocal videos is likewise consistent with the notion of MamL C-terminal interactions with mammalian molecular motors (manuscript submitted).

We further investigated the role of the MamL C-terminal using a Tomato-MamL_trunc_ mutant. When individually expressed in mammalian cells, MamL_trunc_ displays both diffuse and punctate fluorescent patterns, the former more common than the latter. The punctate particles in cells that express MamL_trunc_ are also mobile, suggesting that removal of the C-terminal peptide may affect the affinity of the interaction between MamL and cytoskeletal elements. When Tomato-MamL_trunc_ is co-expressed with EGFP-MamI, three different expression patterns were observed: diffuse MamI and MamL_trunc_ (yellow diffuse), diffuse MamI but punctate MamL_trunc_ (green diffuse but red punctate), or punctate MamI and MamL_trunc_ (yellow punctate). Since colocalization of the two proteins is not always observed, we speculate that the C-terminal of MamL also promotes its interaction with MamI. Possibly, cells displaying green diffuse but red punctate fluorescence represent a transitional state of MamI/MamL_trunc_ interaction where MamI protein is in the process of reorganizing from a net-like to punctate structure. Such a transitional state, however, is not seen in cells co-expressing MamI and full-length MamL.

Whether or not the mobility of Tomato-MamL can be attributed to interactions with eukaryotic molecular motors remains uncertain. Nevertheless, the three major groups of eukaryotic cytoskeletal proteins - actin, tubulin and intermediate filament proteins - are represented in prokaryotes (32) and evolutionarily conserved in MTB (10, 33), as is the magnetosome structure, which has been likened to a bacterial organelle (11). Potentially, MamL has an underlying role in subcellular movement of the magnetosome, known to be linked by MamK (a non-essential magnetosome protein) to actin-like filaments of the bacterial cytoskeleton (34). In support of bacterial-mammalian protein interactions, a functional role for MamM, a magnetosome-associated cation diffusion facilitator (CDF) protein, in modelling structural and regulatory changes in mammalian CDF homologues, has been previously reported (35).

Both MamI and MamL have been implicated in membrane invagination in MTB (36) and thus could both strongly influence membrane curvature and liposome size. A second possible explanation is that, in the absence of MamI, the detected particles strongly associate with larger structures, like the cytoskeleton, resulting in constrained (rather than free) Brownian motion and a very small apparent diffusion coefficient. Alternatively, partial expression of the magnetosome structure may lead to promiscuous interactions with mammalian components (*i.e.,* molecular motors) that would not exist in the presence of the full complement of magnetosome gene products. In any case, the data presented here point to a notable interaction between MamI and MamL, with consequences related to intracellular mobility of the resulting particle. Conceivably, the rudimentary magnetosome structure has built-in motility, in part to facilitate magnetotaxis (37). The nature of additional potential magnetosome protein-protein interactions is currently under investigation (2, 4) and may rely on ionic interactions characteristic of CPP.

We have previously hypothesized that essential magnetosome proteins form a rudimentary structure upon which the membrane-enclosed biomineral is synthesized (4). The data herein support this hypothesis and provide the first confirmation that MamI and MamL interact, even in the complex intracellular compartment of the mammalian cell. This finding is consistent with their proposed role in providing a docking site on the vesicle membrane for additional magnetosome protein interactions. Moreover, the unique patterns of cellular localization observed with MamI and MamL fluorescent fusion proteins suggest that these magnetosome proteins may have additional functionality not previously recognized. Among these is the possibility of creating a scaffold on which a rudimentary magnetosome-like nanoparticle may be assembled for molecular imaging with MRI. Once the relationship between essential magnetosome elements is more fully understood, the regulation of this structure will be possible and permit the selective use of discrete steps in iron biomineralization. This includes regulation of its subcellular localization, timing of assembly or disassembly, and ultimately, control of the size, shape and composition of the crystal. Cellular MRI may be influenced by all these factors, permitting *in vivo* detection of genetically-regulated iron contrast in cells of all kinds, from bacteria (38) to human.

### Experimental procedures

To better understand the nature of magnetosome assembly, we hypothesized that essential magnetosome genes provide the root structure upon which biomineralization depends. To test this theory, we examined the ability of two such proteins to interact in a foreign cell environment. Magnetosome genes *mamI* and *mamL*, as well as a truncated form of *mamL,* were expressed as fluorescent fusion proteins in a mammalian cell line to track their subcellular localization and potential for interaction using confocal microscopy. Magnetosome protein colocalization was examined in cell extracts using co-immunoprecipitation and fluorescence correlation spectroscopy to characterize (changes in) particle diffusion, radius, and intrinsic brightness.

### Molecular Cloning

Magnetosome genes *mamI* and *mamL* were amplified by PCR from the genomic DNA of *Magnetospirillum magneticum* strain AMB-1 (ATCC 700264) using custom primers (Table S1). The *mamI* and *mamL* amplicons were purified using a PCR clean-up kit (Invitrogen, Life Technologies, Burlington, Canada); digested with appropriate restriction enzymes (Table S1); and purified once more, prior to insertion in the molecular cloning vectors pEGFP-C1 (Clontech) and ptdTomato-C1 (Clontech), respectively. After propagation in *Escherichia coli* strain XL10GOLD, the vector-insert plasmid constructs were purified and used for mammalian cell transfection, as outlined below.

To create MamL_trunc_, the last 15 amino acids from the C-terminal of MamL were removed by PCR site-directed mutagenesis. Briefly, primers were designed that included a stop codon before the 15 amino acid peptide. These primers were then used in PCR amplification of the truncated *mamL* gene, which was then inserted into the ptdTomato-C1 vector with restriction enzymes EcoRI and BglII (Table S2).

To refine the co-expression of MamI and MamL, the pSF-EMCV-*FLuc* vector was used to generate FLAG-tagged MamL under puromycin selection. Interaction between magnetosome proteins could then be identified using EGFP-MamI. Primers flanking *mamL* in the ptdTomato construct were designed to include a FLAG tag (DYKDDDDK) for immunodetection (Table S3). The FLAG*-mamL* insert was amplified using PCR, purified using a PCR clean-up kit (Invitrogen, Life Technologies, Burlington, Canada), and digested using SacI and EcoRI (Table S3). FLAG*-mamL* was then inserted into the pSF-EMCV-FLuc vector and propagated in *Escherichia coli* strain XL10GOLD.

### Cell Culture and Transfection

MDA-MB-435 cells (ATCC HTB-129; derived from an adult female and characterized as a melanoma cell line) are a model of aggressive tumorigenesis (22). Cells were cultured in 100 mm polystyrene-coated cell culture dishes (CELLSTAR, VWR International, Mississauga, Canada) with Dulbecco’s Modified Eagle Medium (DMEM) containing 1 g/L glucose (Gibco, Life Technologies, Burlington, Canada), 10% fetal bovine serum (FBS; Gibco), 4 U/mL penicillin, and 4 µg/mL streptomycin at 37°C with 5% CO_2_. To create cell lines expressing the enhanced green fluorescent protein (EGFP)-MamI fusion protein or the red fluorescent protein tdTomato (Tomato)-MamL fusion protein, cells were grown to 60-70% confluency on a 100 mm dish and transfected using Lipofectamine 2000 (Invitrogen), according to company protocol, using 8 µg of pEGFP-*mamI* or ptdTomato-*mamL*, respectively. For co-expression of both pEGFP-*mamI* and ptdTomato-*mamL,* cells stably expressing Tomato-MamL were transfected with 8 µg of pEGFP-*mamI*. After 16 hours transfections were stopped, and cells were placed in full medium for 48 hours before commencing antibiotic selection.

To create cell lines expressing Tomato-MamL_trunc,_ parental MDA-MB-435 cells were transfected with 4 µg of Tomato-*mamL_trunc_* DNA and 12 µg of Lipofectamine 2000. To evaluate whether MamL_trunc_ interacts with MamI, transient transfections were performed. MDA-MB-435/EGFP-MamI (high fluorescence intensity) and MDA-MB-435/EGFP-MamI (medium fluorescence intensity) cells on 35 mm glass-bottom cell culture dishes were each transfected with 1 µg of Tomato-*mamL_trunc_* DNA and 2 µg of Lipofectamine 2000.

To create cell lines co-expressing EGFP-MamI and FLAG-MamL, MDA-MB-435/EGFP-MamI cells were grown to 60-70% confluency on a 100 mm dish and transfected using Lipofectamine 2000 (Invitrogen), according to company protocol, using 8 µg of pSF-FLAG-*mamL*-EMCV-FLuc. After 16 hours transfection was stopped, and cells were placed in full medium for 48 hours before commencing antibiotic selection.

### Selection of Stable Cell Lines

To select cells stably expressing EGFP-MamI and/or Tomato-MamL fusion proteins, transfected cells were grown in the presence of 500 μg/mL geneticin (G418; Gibco). To enrich the populations of fluorescing cells, we used fluorescence activated cell sorting (FACS; Robarts Research Institute, London, Canada). Two cell populations, one each with high and medium fluorescent intensities, were obtained for western blots (only high) and confocal (only high) fluorescence microscopy, or FCS (only medium), respectively.

To select cells stably expressing Tomato-MamL_trunc_, transfected cells were grown in the presence of 500 μg/mL geneticin (G418; Gibco).

To select cells stably co-expressing EGFP-MamI and FLAG-MamL, transfected cells were grown in the presence of 500 μg/mL geneticin (G418; Gibco) and 500 ng/mL puromycin (Gibco).

### Protein Sample Preparation

Stably transfected cells were cultured to 70% confluency on a 100 mm dish, then washed twice using 10 mL phosphate buffered saline pH 7.4 (PBS, 137 mM NaCl/2.7 mM KCl/10 mM Na_2_HPO_4_). Four to five dishes of cells were then collected into a 1 mL lysis solution containing 850 μL of radioimmunoprecipitation assay buffer (RIPA, 10 mM Tris-HCl pH 7.5/140 mM NaCl/1% NP-40/1% sodium deoxycholate/0.1% sodium dodecyl sulfate [SDS]) and 150 μL of Complete Mini protease inhibitor cocktail (Roche Diagnostic Systems, Laval, Canada). Cells were then sonicated using three 12-second bursts of a Sonic Dismembrator (model 500, Thermo Fischer Scientific, Ottawa, Canada) at an amplitude of 30%. Total amount of protein was quantified using the BCA assay (39).

### Western Blot

Protein samples of MDA-MB-435 cells stably expressing EGFP (40 µg), Tomato (40 µg), EGFP-MamI (20 µg), or Tomato-MamL (20 µg), or stably co-expressing Tomato-MamL/EGFP-MamI (80 µg) were reduced with 100 mM dithiothreitol (DTT) in sample preparation buffer (1 M Tris-HCl pH 6.8/10% SDS/0.1% Bromophenol Blue/43% glycerol) and heated at 85°C for at least 5 min. Reduced samples were then subjected to discontinuous SDS polyacrylamide gel electrophoresis (SDS-PAGE) using a 10% running gel (14% running gel for FLAG-MamL detection). Protein was transferred onto a nitrocellulose blot using the Original iBlot Gel Transfer Device (Life Technologies, Burlington, Canada).

For EGFP detection, nonspecific protein binding was blocked in 5% bovine serum albumin (BSA)/Tris-buffered saline pH 7.4 (TBS) for 3 h at room temperature. Blots were then incubated for 15 h in 1:1000 mouse α-GFP (Invitrogen)/3% BSA/TBS/0.02 % sodium azide (TBSA); then washed using TBS/0.1% Tween 20 (TBST; Sigma-Aldrich, Oakville, Canada) for 30 min with 4 changes of buffer; and incubated for 2 h in 1:20,000 horseradish peroxidase (HRP)-conjugated goat α-mouse IgG (Sigma-Aldrich)/1% BSA/TBS. All incubations were performed at room temperature. Blots were then washed with 0.1% TBST for 30 min with 4 changes of buffer and imaged using the Chemigenius Gel Doc (Syngene). A chemiluminescent signal was detected using SuperSignal West Pico Chemiluminescent Substrate (Thermo Fischer Scientific), according to the manufacturer’s instructions.

For Tomato detection, blots were blocked in 3% BSA/TBSA for approximately 18 h at room temperature and then incubated for 18 h in 1:1000 primary goat α-tdTomato (MyBioSource, San Diego, USA)/3% BSA/TBSA at 4°C. After washing in 0.1% TBST as described above, blots were incubated for 1 h in 1:20,000 HRP-conjugated rabbit α-goat IgG (Sigma-Aldrich)/1% BSA/TBS at room temperature.

For FLAG detection, blots were blocked in 3% BSA/TBS overnight and then incubated for 18 h in 1:1000 primary mouse α-FLAG (Thermo Fisher Scientific)/3% BSA/TBSA. After washing in 0.1% TBST as described above, blots were incubated for 2 h in 1:20,000 HRP-conjugated goat α-mouse IgG (Sigma-Aldrich)/1% BSA/TBS. All incubations were done at room temperature.

Glyceraldehyde 3-phosphate dehydrogenase (GAPDH) was used as a loading control. For GAPDH detection, blots were placed in stripping solution (1 M Tris-HCl pH 6.8/10% SDS/0.016% β-mercaptoethanol) and agitated in a 37°C water bath for 30 min prior to washing in 0.1% TBST and blocking in 5% BSA/TBS. The primary and secondary antibodies were 1:2000 rabbit α-GAPDH (Sigma-Aldrich)/3% BSA/TBSA and 1:20,000 HRP-conjugated goat α-rabbit IgG (Sigma-Aldrich)/1% BSA/TBS, respectively.

### Co-immunoprecipitation

Protein samples were prepared and quantified as described above. For immunoprecipitation, 5 mg of GFP-MamI or FLAG-MamL total cell extract was mixed with 5 µL of mouse α-GFP and incubated for 1h at 4°C. Protein-G Sepharose beads (Cytiva, Vancouver, BC, Canada) were washed and added to each protein-antibody mixture and then incubated 1h at 4°C. Unbound protein was washed away with RIPA buffer and 10 mM Tris-HCl. Protein associated with Protein-G Sepharose was resuspended in 40 uL of sample buffer (125 mM Tris-HCl pH 6.8/3.3% SDS/100 mM DTT) and heated to 80°C for 15min to dissociate protein from beads. Glycerol (10% v/v) and bromophenol blue (∼0.1% w/v) were added to each sample prior to storing at −20°C until SDS-PAGE.

The co-immunoprecipitation blot was blocked in 4% BSA/TBSA for approximately 5 h then incubated for 18 h in 1:2000 primary goat α-FLAG (Thermo Fisher Scientific)/3% BSA/TBSA at room temperature. After washing in 0.1% TBST as described above, blots were incubated for 1 h in 1:20,000 HRP-conjugated rabbit α-goat IgG (Sigma-Aldrich)/1% BSA/TBS.

### Confocal Imaging

Stably transfected cell lines were examined with confocal fluorescence microscopy (Nikon A1R Confocal Microscope) to confirm expression and characterize the intracellular localization of EGFP-MamI and/or Tomato-MamL fusion proteins. In preparation for confocal microscopy, cells were cultured in a 35 mm glass-bottom dish (MatTek Corporation, Cedarlane, Burlington, Canada) for 48 hours. On the day of imaging, the dish was placed in a stage-top incubator to maintain 37°C and 5% CO_2_. Images and cines were captured using a Galvano scanner with NIS-Elements AR 5.11.01 (Nikon Instruments Inc.), using a 20X objective with 0.75 numerical aperture. To capture images of cells expressing a single fluorophore, the FITC microscope filter (495 nm excitation/519 nm emission) was used for cells expressing the EGFP fluorophore and the TRITC microscope filter (557 nm excitation/576 nm emission) was used for cells expressing the Tomato fluorophore. To capture images of cells co-expressing both fluorophores (EGFP and Tomato), the FITC and TRITC filters were turned on simultaneously. Captured images of cells in both channels were then merged in Adobe Photoshop CS6.

Cines were acquired with the time lapse function in NIS-Elements AR 5.11.01, which acquires a picture every 2 s for a total of 60 s. Cines were captured in either channel or both channels simultaneously, as described above. The NIS-Elements software automatically generated a time lapse video with single or merged channels. This video was then edited in Adobe Photoshop CS6 and exported as a GIF file.

### Fluorescence Correlation Spectroscopy (FCS)

Lysed samples of parental MDA-MB-435, MDA-MB-435/EGFP-MamI, MDA-MB-435/Tomato-MamL, and MDA-MB-435/Tomato-MamL/EGFP-MamI cells were collected following the protein sample preparation protocol described above. These samples were stored at −20^°^C until FCS data was acquired using an Evotec Insight confocal instrument (Evotech Technologies, Hamburg, Germany, now Perkin-Elmer, Waltham, USA) equipped with a 40X water immersion objective (Olympus, Tokyo, Japan) used in combination with a 40 μm confocal pinhole. A 488 nm excitation source (power 20 μW) was used for samples containing EGFP and a 532 nm excitation source (power 10 μW) was used for samples containing Tomato. A calibration step was first performed with fluorescent dyes to determine the dimensions of the detection volumes (in both EGFP and Tomato channels). The viscosity and background fluorescence of the cell extract was obtained through lysis of the parental MDA-MB-435 cells.

Solubilized samples of total cellular protein (2 mg/ml) were then loaded into the wells of a glass-bottom 96-well plate (Greiner Sensoplate, Sigma-Aldrich) and FCS measurements acquired for each sample (3 to 5 repeats of 20 to 60 s measurements, acquired approximately 5 µm above the glass coverslip to avoid mismatch in the refractive index between immersion water and cell extract).

Both autocorrelation functions and photon counting histograms were generated and analyzed, using either a one-component or two-component model, and taking into account the photophysics of the fluorescent proteins. The diffusion coefficient (*D*) and intrinsic brightness (*B*) of the fluorescent particles detected in each sample were then calculated according to standard protocols (40). The apparent radius of the fluorescent species detected (*R*) was calculated using the Stokes-Einstein relationship: *D = kT/*(6*πηR*), where *k* is Boltzmann’s constant, *T* = 300 K is the absolute temperature, and viscosity of the cell extract, *η* = 1.29 *η_water_* = 1.15 x 10^-3^ Pa.s, was measured in a separate experiment.

### Statistical Analysis

All statistical tests were performed using GraphPad Prism version 8. An unpaired t-test was used to evaluate statistically significant differences (p < 0.05) between the velocities of Tomato-MamL and Tomato-MamL/EGFP-MamI particles. An unpaired t-test was also used to identify any significant difference between the FCS intrinsic brightness of Tomato and Tomato-MamL particles and of EGFP and EGFP-MamI particles. Significant differences between the FCS parameters of EGFP, EGFP-MamI, and EGFP-MamI/Tomato-MamL structures were determined using one-way analysis of variance (ANOVA). Similarly, significant differences between FCS parameters of Tomato, Tomato-MamL, and Tomato-MamL/EGFP-MamI structures were determined using one-way ANOVA.

## Data availability

All data needed to evaluate the conclusions in the paper are present in the paper and/or the Supporting Information.

## Acknowledgements

The authors thank Dr. Kevin P. White, Dr. Jeremy P. Burton, and Dr. Shoogo Ueno for providing critical feedback on the manuscript.

## Author Contributions

Sun Q contributed to experimental design, data collection and analysis, creation of figures and tables, manuscript writing, editing, and revising. Yu L contributed to data collection and analysis and creation of figures and tables. Donnelly SC and Fradin C contributed to experimental design, data collection and analysis, creation of figures and tables, and manuscript writing, editing, and revising. Thompson RT contributed to experimental funding and manuscript revising. Prato FS contributed to experimental funding and to manuscript editing and revising. Goldhawk DE contributed to experimental design and manuscript writing, editing, and revising.

### Funding and additional information

This research was supported by the Ontario Research Fund (RE07-021) in partnership with Multi-Magnetics Inc.; by the Cancer Imaging Network of Ontario through Cancer Care Ontario; and by Discovery Grants (2014–05589, 2020–04125) from the Natural Sciences and Engineering Research Council of Canada.

## Supporting Information

### List of Supporting Information

**Table S1.**
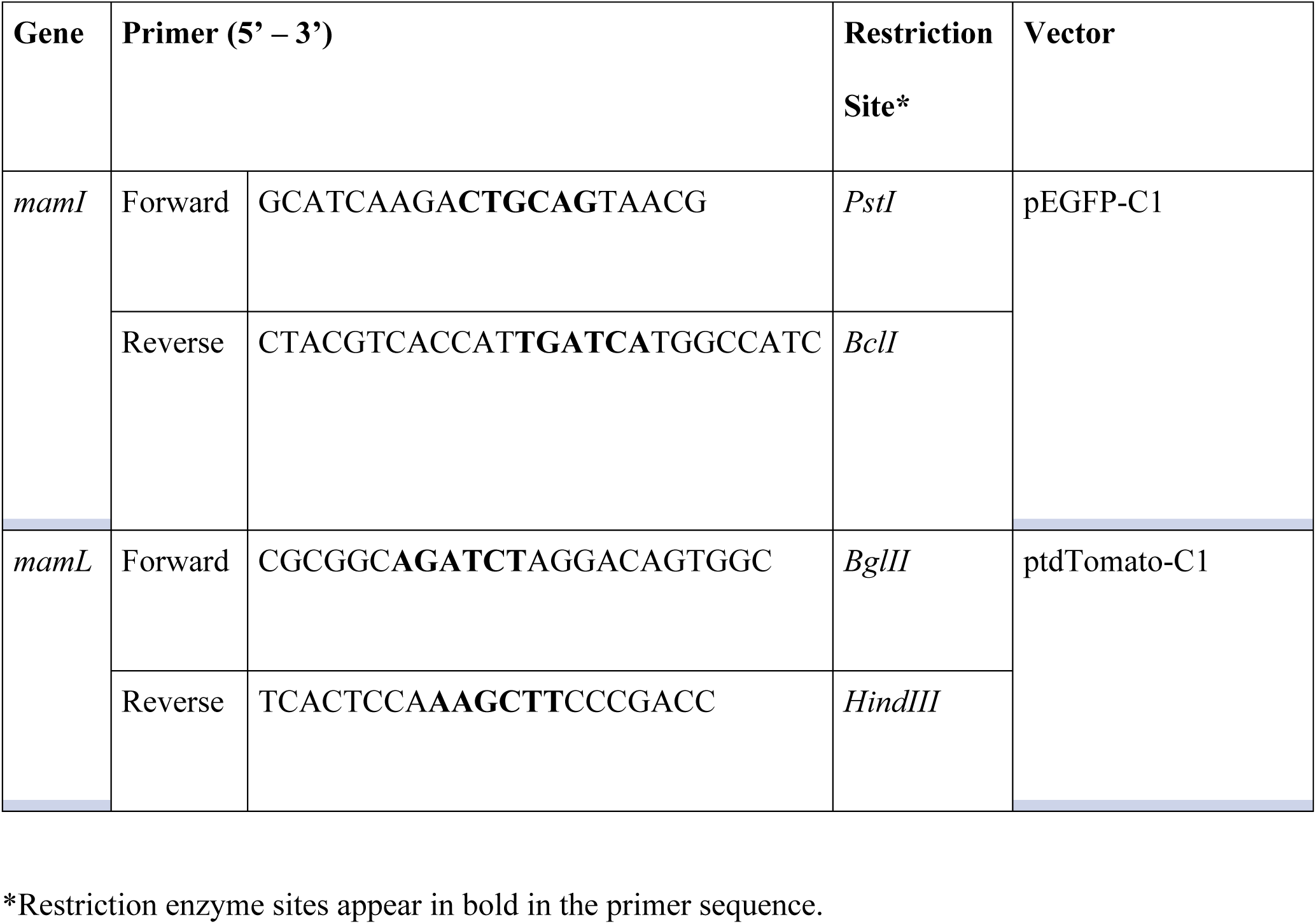
Primer design for the cloning of MTB genes mamI and mamL.

**Table S2.**
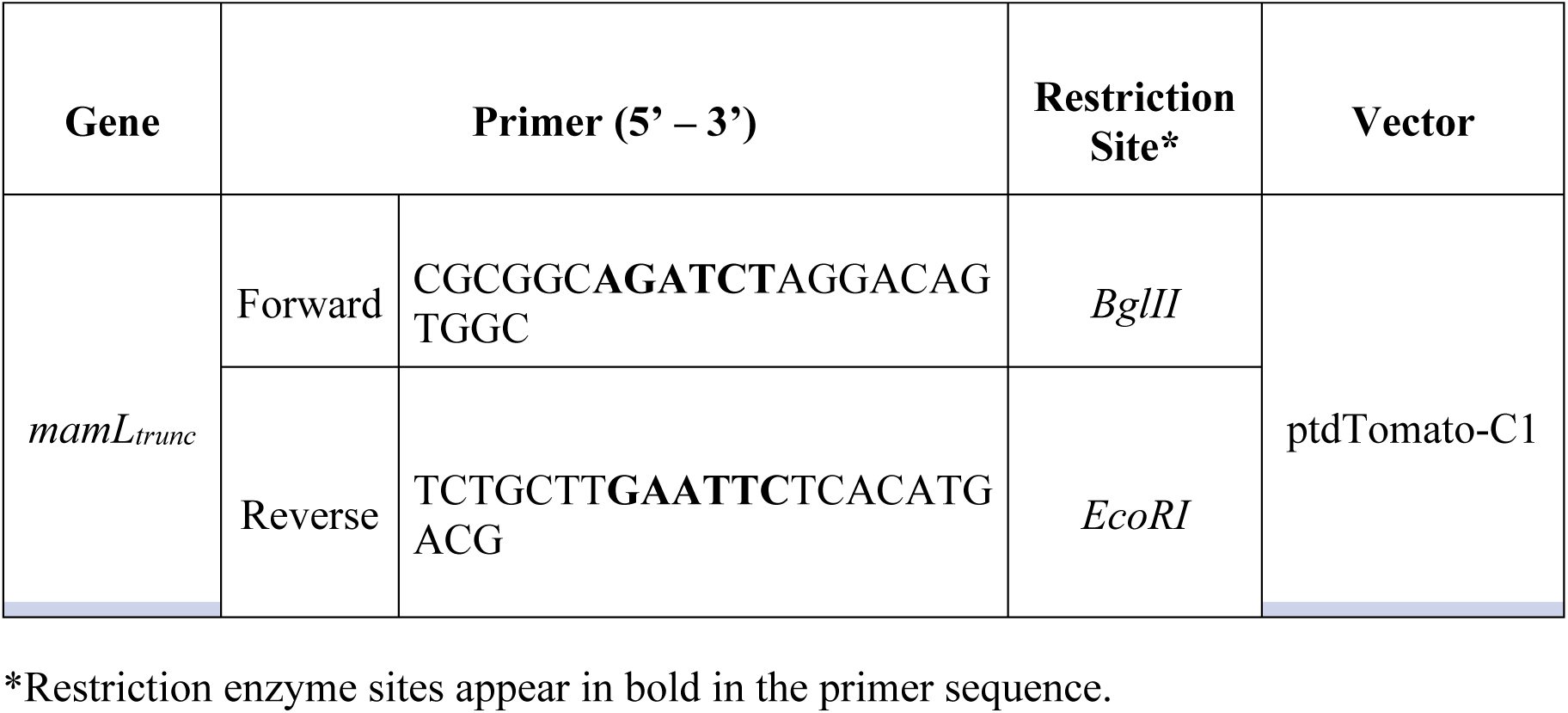
Primer design for the cloning of MTB gene *mamL_trunc_* into the ptdTomato-C1 vector.

**Table S3.**
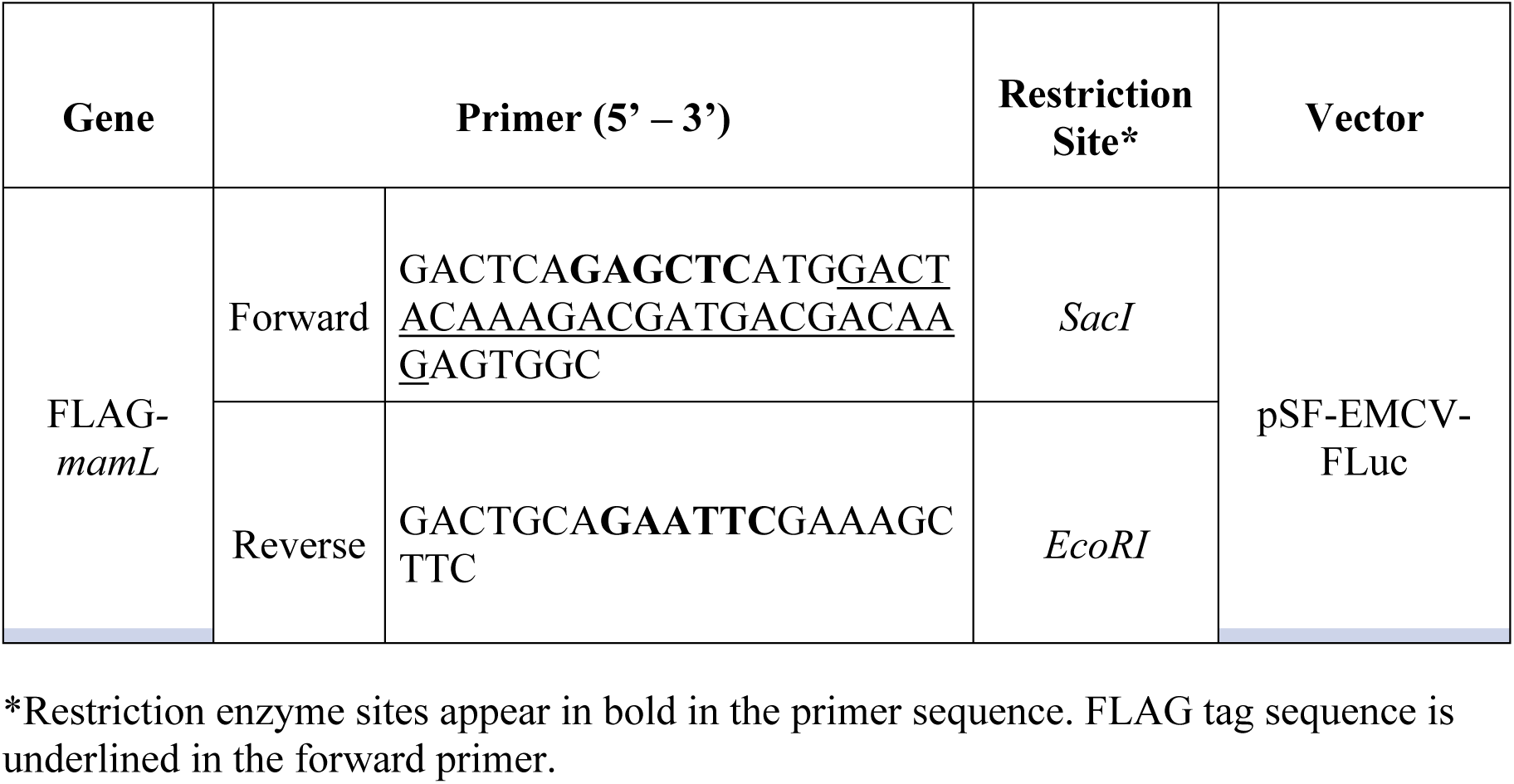
Primer design for the cloning of FLAG-*mamL* into the pSF-EMCV-FLuc vector.

**Figure S1.**
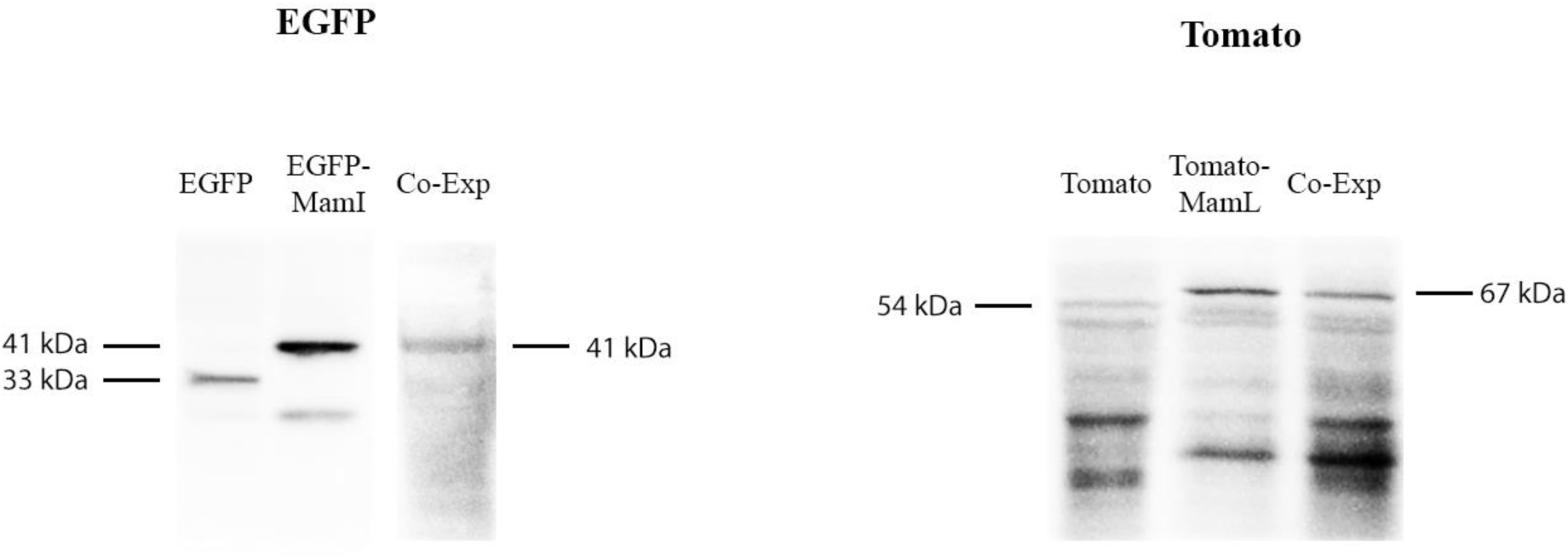
Immunoblots of total cellular protein from mammalian cells. Total cellular protein from MDA-MB-435 cells stably expressing EGFP, EGFP-MamI, and co-expressing EGFP-MamI/Tomato-MamL were examined with mouse α-EGFP (left). Total cellular protein from cells stably expressing Tomato, Tomato-MamL, and co-expressing EGFP-MamI/Tomato-MamL were examined with α-Tomato (right). Full-length western blots are displayed.

**Figure S2.**
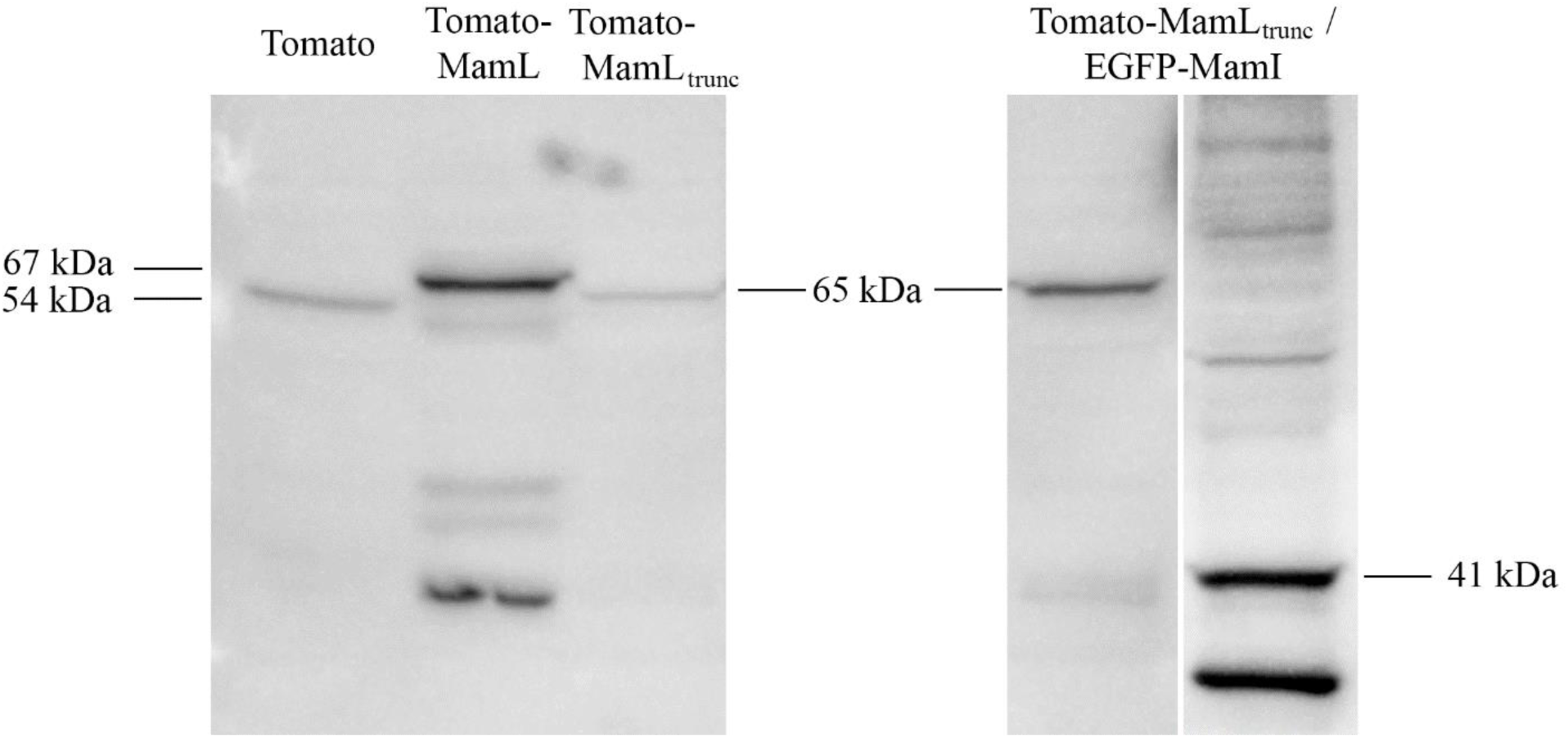
Immunoblots of total cellular protein from mammalian cells. Total cellular protein from MDA-MB-435 cells stably expressing Tomato, Tomato-MamL, Tomato-MamL_trunc_, and co-expressing Tomato-MamL_trunc_/EGFP-MamI were examined with rabbit α-Tomato or mouse α-EGFP (far right lane). Full-length western blots are displayed.

**Figure S3.**
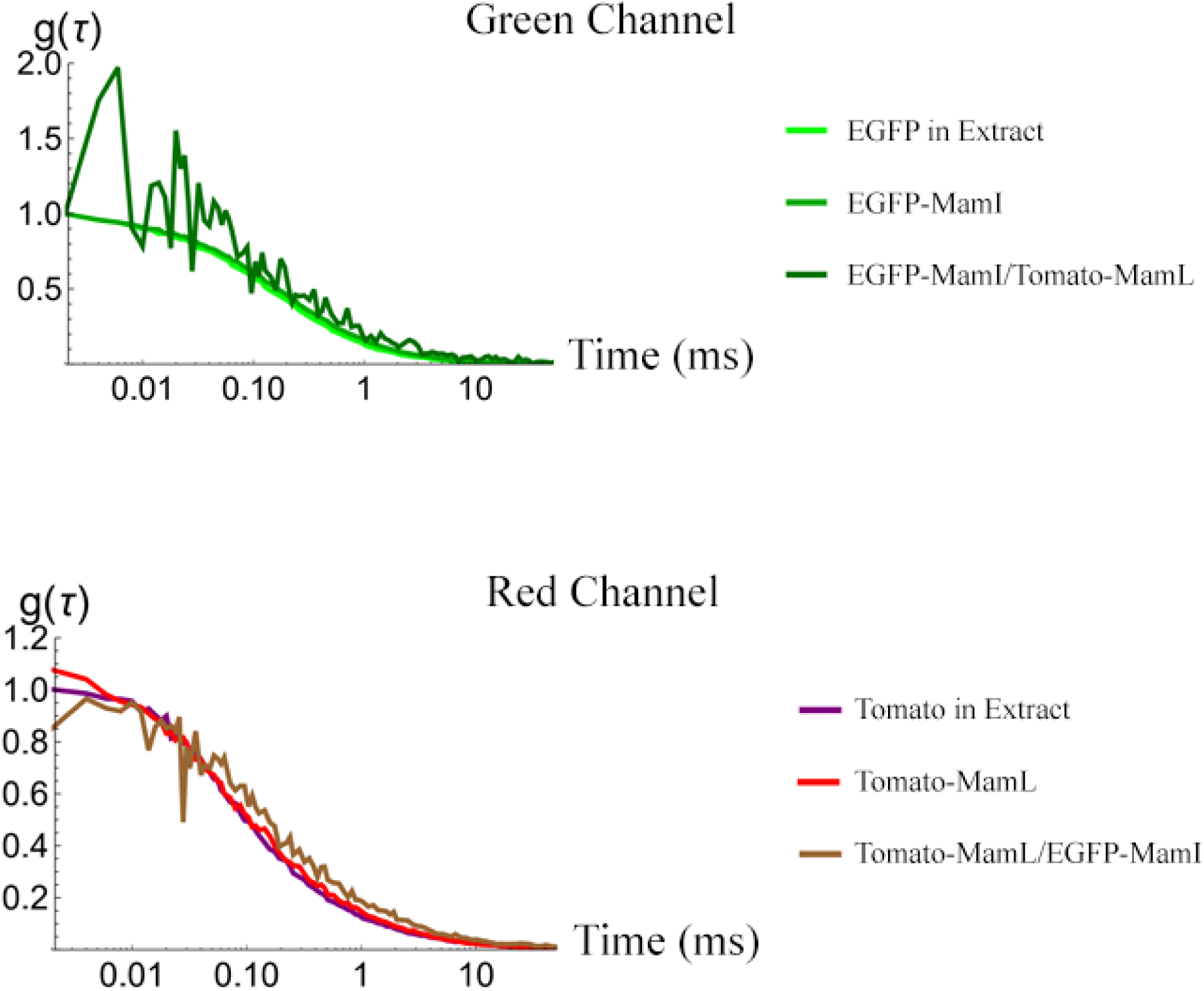
Autocorrelation curves obtained from fluorescence correlation spectroscopy of mammalian cells. Curves in the green channel (top) are signals collected from EGFP (light green), EGFP-MamI (green), or EGFP-MamI/Tomato-MamL (dark green) in lysed samples containing total cellular protein. Curves in the red channel (bottom) are signals collected from Tomato (purple), Tomato-MamL (red), or Tomato-MamL/EGFP-MamI (brown). Curves are semi-log plots with autocorrelation function (g(τ)) on the y-axis and lag time (ms) on the x-axis.

**Figure S4.**
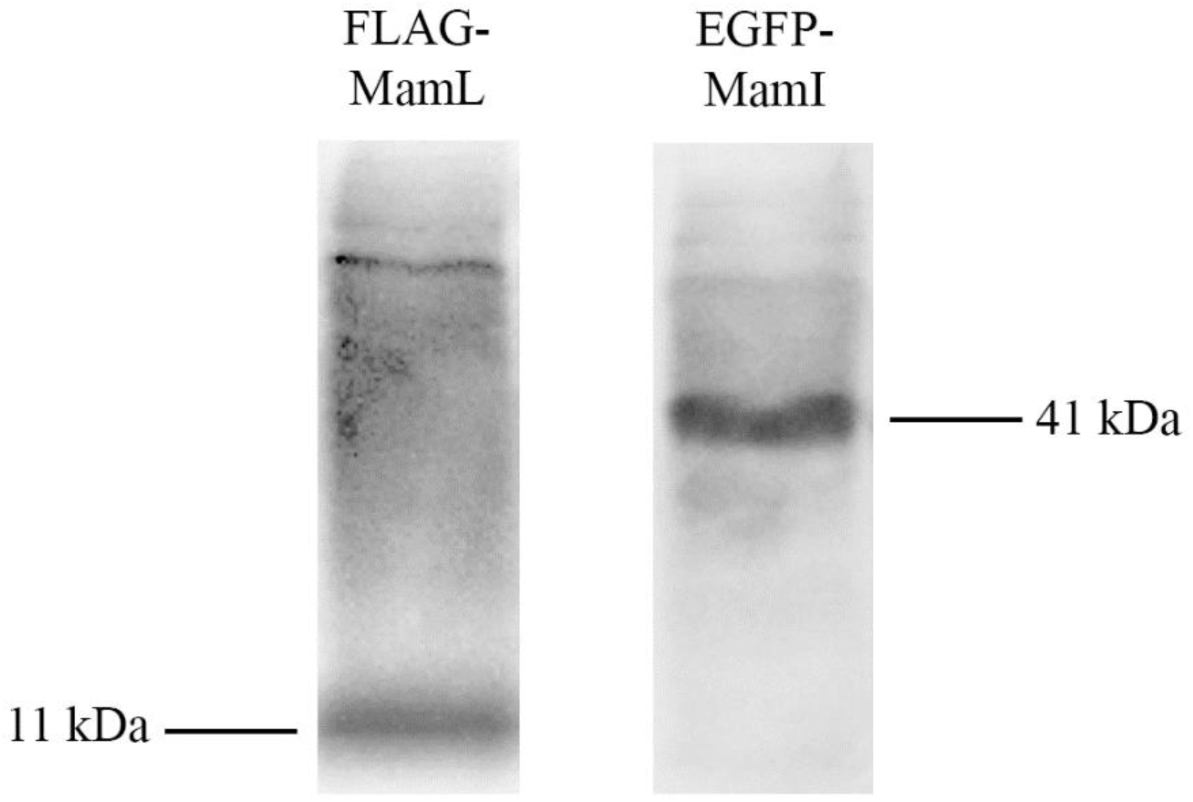
Immunoblots of total cellular protein from mammalian cells. Total cellular protein from MDA-MB-435 cells stably expressing FLAG-MamL was examined with mouse α-FLAG (left) or stably expressing EGFP-MamI was examined with mouse α-EGFP (right). Full-length western blots are displayed.

(GIF Attachment)

**Movie 1. Time lapse of mammalian cells stably expressing each fluorescent magnetosome fusion protein.** Cines were captured on a Nikon A1R confocal microscope at a rate of 1 frame every 2 seconds for a total of 60 seconds. (**A**) A representative time lapse of EGFP-MamI (from the cell in **Figure 2C**) shows a net-like pattern of fluorescence. The blue arrow points to a junction in this pattern. (**B**) A representative time lapse of Tomato-MamL (from the cell in **Figure 3C**) displays punctate fluorescence, with the blue arrow following one such structure to track its path in each frame.

(GIF Attachment)

**Movie 2. Time lapse of mammalian cells stably co-expressing fluorescent magnetosome fusion proteins.** Cines were captured on a Nikon A1R confocal microscope at a rate of 1 frame every 2 seconds for a total of 60 seconds. A representative time lapse of Tomato-MamL_trunc_ (from the cell in **Figure 5A**) shows a punctate, mobile pattern of fluorescence. The blue arrow follows a punctate structure to track its path in each frame.

(GIF Attachment)

**Movie 3. Time lapse of mammalian cells stably co-expressing fluorescent magnetosome fusion proteins.** Each cine was independently captured on a Nikon A1R confocal microscope at a rate of 1 frame every 2 seconds for a total of 60 seconds. (**A**) A representative time lapse, acquired from the FITC channel, shows EGFP-MamI (from the cell in **Figure 6B**) adopting a mobile, punctate fluorescence pattern when co-expressed with Tomato-MamL. (**B**) A representative time lapse, acquired from the TRITC channel, shows how Tomato-MamL (from the cell in **Figure 6C**) retains its mobile, punctate fluorescence pattern when co-expressed with EGFP-MamI. (**C**) A representative time lapse, acquired simultaneously from both FITC and TRITC channels, shows fluorescent fusion protein co-localization (in yellow) as a mobile punctate pattern. The blue arrows in each cine track punctate structures from frame to frame, highlighting their various paths of movement.

(GIF Attachment)

**Movie 4. Time lapse of mammalian cells stably co-expressing fluorescent magnetosome fusion proteins.** Cines were captured on a Nikon A1R confocal microscope at a rate of 1 frame every 2 seconds for a total of 60 seconds. A representative time lapse of EGFP-MamI/Tomato-MamL_trunc_ (from the cell in **Figure 7C**) shows a punctate, mobile pattern of fluorescence (yellow). The blue arrow follows a punctate structure to track its path in each frame.

(GIF Attachment)

**Movie 5. Time lapse of mammalian cells stably co-expressing fluorescent magnetosome fusion proteins.** Cines were captured on a Nikon A1R confocal microscope at a rate of 1 frame every 2 seconds for a total of 60 seconds. A representative time lapse of EGFP-MamI/Tomato-MamL_trunc_ (from the cell in **Figure 7B**) shows a net-like pattern of fluorescence (green) and punctate, mobile pattern of fluorescence (red). The blue arrow follows a red punctate structure to track its path in each frame.

(GIF Attachment)

**Movie 6. Time lapse of mammalian cells stably co-expressing fluorescent magnetosome fusion proteins.** Cines were captured on a Nikon A1R confocal microscope at a rate of 1 frame every 2 seconds for a total of 60 seconds. A representative time lapse of EGFP-MamI/FLAG-MamL (from the cell in **Figure 8**) shows a punctate, mobile pattern of fluorescence (green). The blue arrow follows a punctate structure to track its path in each frame.

