## Supplementary figures and images for "Essential magnetosome proteins MamI and MamL from magnetotactic bacteria interact in mammalian cells"

### Movie 1

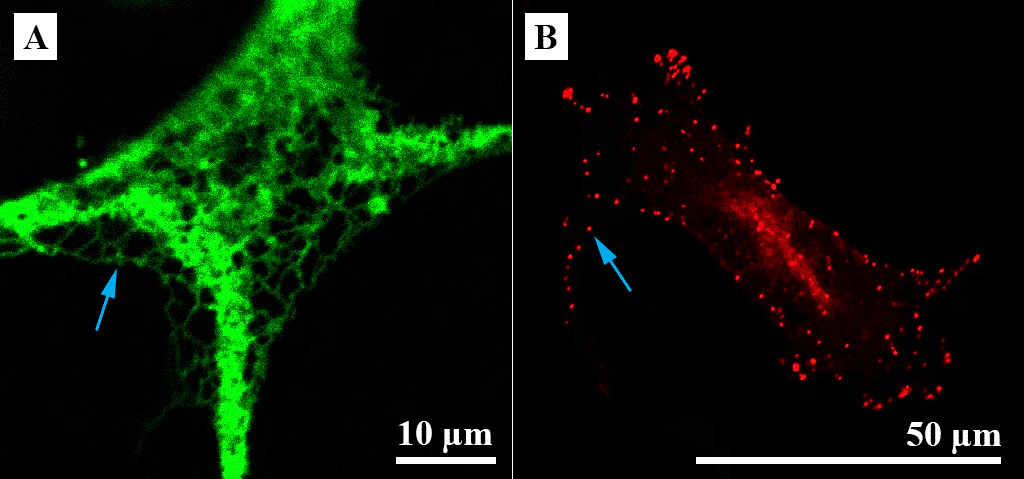

### Movie 2

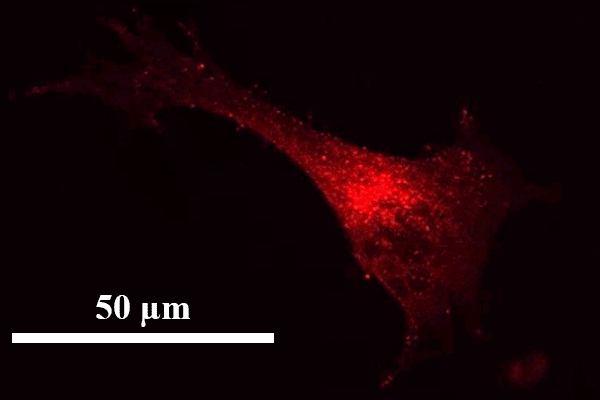

### Movie 3

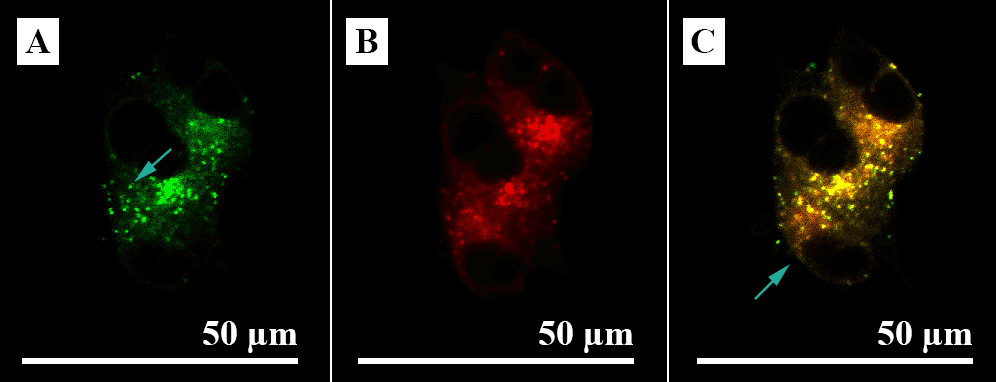

### Movie 4

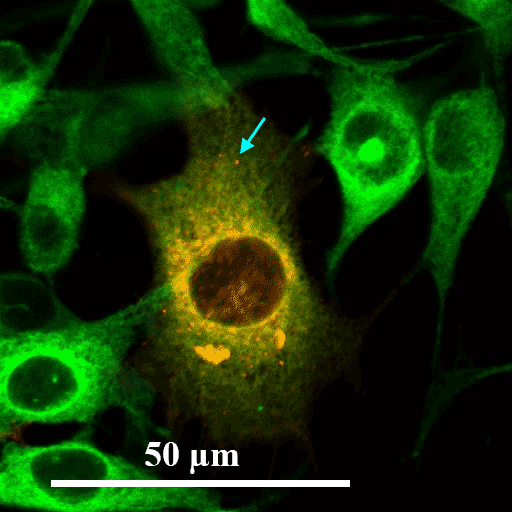

### Movie 5

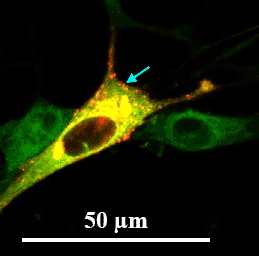

### Movie 6

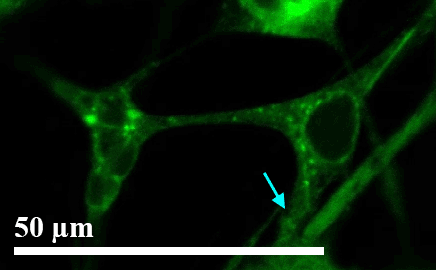
